# Shared Transcriptomic Responses to Distinct Sleep-Wake Manipulations Reveals Multiple Homeostatic Pathways in *Drosophila*

**DOI:** 10.64898/2026.02.28.708752

**Authors:** Clark Rosensweig, Aadish Shah, Shiju Sisobhan, Tomas Andreani, Ravi Allada

## Abstract

Sleep is governed by two processes: a circadian process that times sleep and wake and a homeostatic process that drives sleep as a function of prior wake history. Discovered in the fruit fly *Drosophila*, the *period* (*per*) gene is a “universal” cornerstone of the circadian clock, robustly oscillating at the transcript level, in all organs and tissues and in essentially all animals. We hypothesized that there may be a comparable factor (we term “sleeper”) for sleep homeostasis. To identify sleeper genes, we performed a wide-ranging transcriptomic analysis to identify genes whose expression tracks sleep-wake history in the *Drosophila* brain. We analyzed a variety of methods to manipulate sleep-wake and subsequent rebound including mechanical, thermogenetic, optogenetic, pharmacological and baseline sleep across 7 datasets. Using a log2 fold change threshold of 1, we did not identify any gene that was sleep-wake dependent across all datasets, raising the possibility that genes identified with a single method of sleep manipulation are related to the nature of the manipulation and not necessarily to sleep. Nonetheless, we did identify genes whose expression changed with sleep-wake in a consistent direction across at least 2 datasets. These analyses highlight previous processes implicated in sleep homeostasis such as mitochondrial oxidative phosphorylation as well as provide potentially novel pathways such as ribosome biogenesis. In addition, we also examined genes whose expression correlates with prior sleep-wake history and/or predicted subsequent sleep rebound across these datasets. Among significantly correlated genes we observed some that were correlated with recent (<3h) sleep history while others were correlated with much longer (>6h) time frames suggesting temporally distinct pathways for integrating waking experience. This analysis also highlights specific GO pathways for sleep/wake/rebound. Applying a novel GPT based paper search algorithm, we call *fl.ai*, we identified several sleep-wake regulated genes with demonstrated function in sleep regulation consistent with an *in vivo* homeostatic function. These include those involved in immunity, neuropeptide signaling, glucose transport, and neuronal excitability. Collectively, these studies suggest that sleep homeostasis is a much more molecularly distributed process than the core circadian clock.

## Introduction

Fundamental to an understanding of why we sleep is the homeostatic regulation of sleep. Loss of a night of sleep is followed by a sleep rebound, i.e., an increase in sleep duration or intensity, to recover lost sleep and return to its homeostatic set point. Homeostatic processes are classically modeled as closed loop negative feedback control systems. In the case of sleep, wakefulness is thought to result in the production of factors that steadily accumulate until they hit a threshold when sleep is triggered and the factors return to their set point levels. The circadian clock is thought to time the sleep-wake cycle by altering this homeostatic set point, ensuring consolidated sleep and wake times^1–3^. Yet our molecular understanding of this process is limited, especially related to other fundamental homeostatic processes. Synaptic strength^4, 5^, protein phosphorylation^6–10^, and metabolic processes such as mitochondrial oxidative phosphorylation^11, 12^ and adenosine production^13^, are among the processes that have been proposed as key sleep homeostatic mechanisms.

To address this question, we have been leveraging the fruit fly *Drosophila*. The fly exhibits many of the cardinal properties of sleep including behavioral quiescence, elevated arousal threshold, and homeostatic regulation(reviewed in^14, 15^). *Drosophila* sleep is also important for long term memory consolidation^16^. Moreover, it exhibits electrical correlates and is similarly affected by drugs that also alter sleep in humans, such as the GABA-A agonist gaboxadol (4,5,6,7-tetrahydroisoxazolo-[5,4-c]pyridin-3-ol; THIP)^17^.

Sleep is also governed by a circadian clock. Molecular genetics in *Drosophila* has been pivotal to revealing the “universal” transcriptional feedback loops at the core of circadian timing^18^. CLOCK(CLK) along with its heterodimeric partner CYCLE activates the transcription of *period (per)* and *timeless (tim)*. PER dimerizes with TIM and feeds back to repress CLK transcriptional activation. This results in high amplitude transcript rhythms of *per* and *tim*, among other core clock genes. These feedback loops are evident and largely synchronous across tissues and cell-types including in the brain, such that they are evident even in mixtures of bulk tissues, e.g., the *Drosophila* head^19–27^ or brain^28^. Many of the molecular components and the feedback logic are conserved across essentially all animals.

Inspired by the circadian clock, we reasoned that sleep homeostasis is similarly encoded at the transcriptional level and that it would be evident in whole brains. Many studies have used various methods to deprive sleep and assess the transcriptomic impact^29–39^. Yet these studies do not necessarily distinguish gene expression changes related to the specific method of deprivation from those related to sleep loss. Thus, we reasoned that identifying genes whose expression changes across independent sleep deprivation (SD) methods would allow the discovery of “true” core sleep homeostatic factors. Here we find that overlap between sleep-wake regulated genes using diverse SD methods, while not uniform, is significantly greater than by chance and identifies both established and novel pathways implicated in sleep homeostasis. Moreover, many of these genes have established *in vivo* sleep functions consistent with a *bona fide* role in sleep homeostasis. This work highlights not a single dedicated pathway, as in the core circadian clock, but multiple diverse molecular pathways that each appear to contribute to the homeostatic regulation of sleep.

## Results

To identify key mediators of sleep homeostasis, we reasoned that such factors should be evident independent of the method used to manipulate the sleep-wake cycle. Here we examine flies under baseline sleep-wake conditions (Zeitgeber Time 0,4,8,12,16,20 where ZT0 is lights-on and ZT12 is lights-off), after mechanical SD and after thermogenetically induced sleep and wakefulness (Fig. 1). We applied mechanical SD for 3, 6, and 12 hours ending at ZT0 and each showed significant sleep loss and rebound (Fig. 1C,D). In addition, we induced sleep thermogenetically using R85C10-GAL4 in combination with TrpA1, a temperature-activated cation channel. We observed significant sleep promotion after 12h of daytime activation (29°C) and significant subsequent anti-rebound, consistent with the idea that sleep drive was dissipated (Fig. 2). Whole brains were dissected after each manipulation and then subjected to transcriptomic analysis using RNA-seq. In addition to the datasets we collected, we also analyzed published RNA-seq datasets collected from fly brains after sleep induction via optogenetic activation of dorsal fan shaped body and ventral nerve cord “Bowtie” neurons (R23E10^40^) using the red light-activated ion channel ChRimson (ChR) or chemical induction using THIP^41^. To assess the validity of our technique, we examined the expression of canonical clock genes (*per*, *tim*, and *Clk*) and observed high amplitude oscillations ranging from 3-10 fold in magnitude (Fig. 3) with the expected phase^28, 33, 42, 43^. Importantly, these data demonstrate that we can observe high amplitude transcript oscillations using whole brain samples.

**Figure 1.**
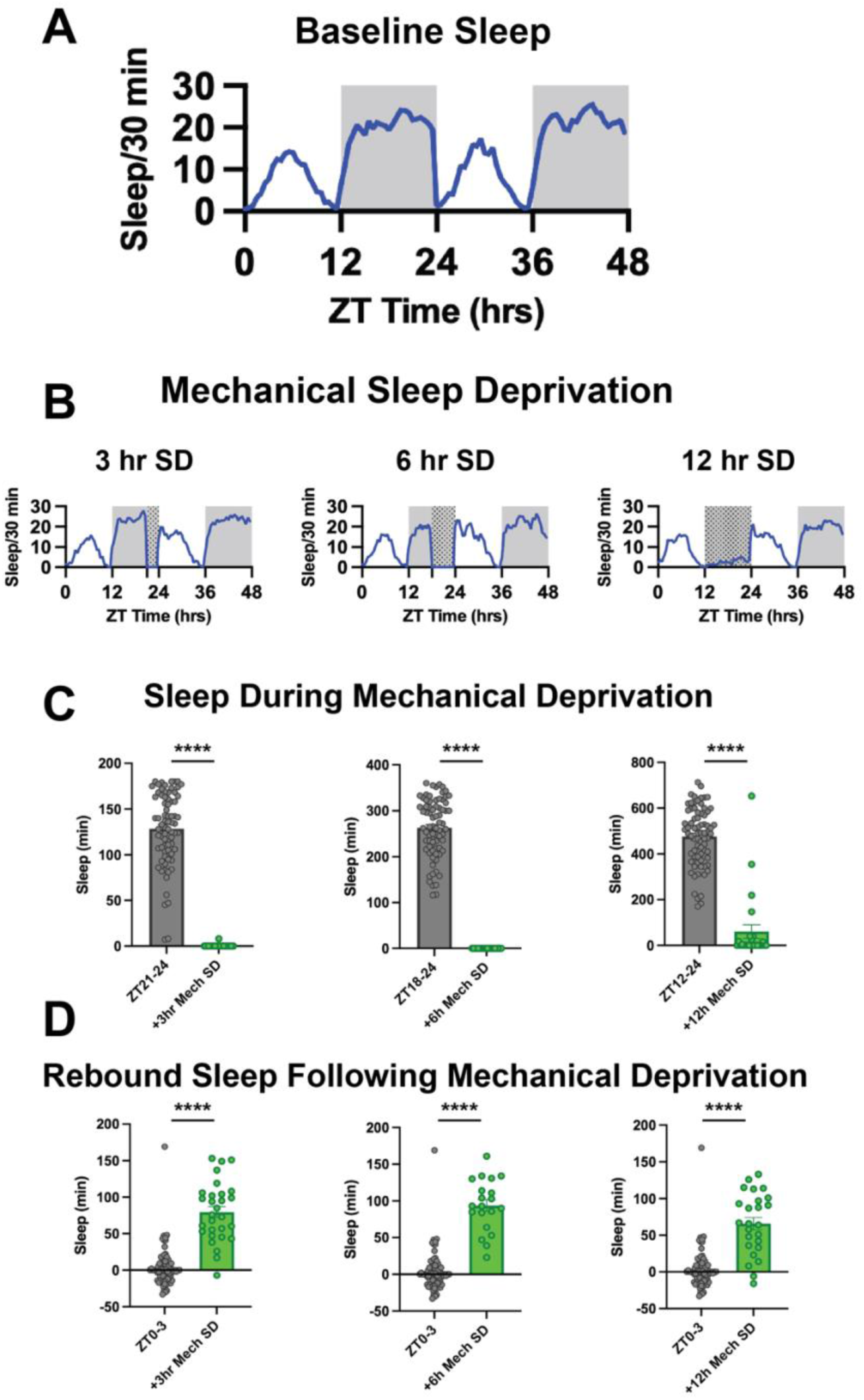
Sleep Under Baseline and Mechanical Sleep Deprivation. A. Minutes of sleep per 30 min interval over two circadian cycles in 12:12 hr LD conditions from *iso31* wild-type flies (n = 74). B.Minutes of sleep per 30 min interval over two circadian cycles for *iso31* wild-type flies across three conditions. The three panels show flies that received a 3 hr (n = 29), 6 hr (n = 20), and 12 hr (n = 25) mechanical sleep deprivation at the end of the first dark period respectively. C. Quantitation of the data shown in panels A and B. Each SD condition is compared to the equivalent time interval during the first night of the circadian data in panel A. Total minutes of sleep during each interval are shown for each fly. Control flies (n = 74), 3 hr SD (n = 29), 6 hr SD (n = 20), and 12 hr SD (n = 25). **** p<0.0001 by Welch’s t-test. D. Quantitation of the rebound shown in panels A and B. Graphs show the total sleep in the first 3 hrs after sleep deprivation. Each SD condition is compared to the equivalent time interval (ZT0-3) on the second day of the circadian data in panel A. Total minutes of sleep during each interval are shown for each fly. Control flies (n = 74), 3 hr SD (n = 29), 6 hr SD (n = 20), and 12 hr SD (n = 25). **** p<0.0001 by Welch’s t-test.

**Figure 2.**
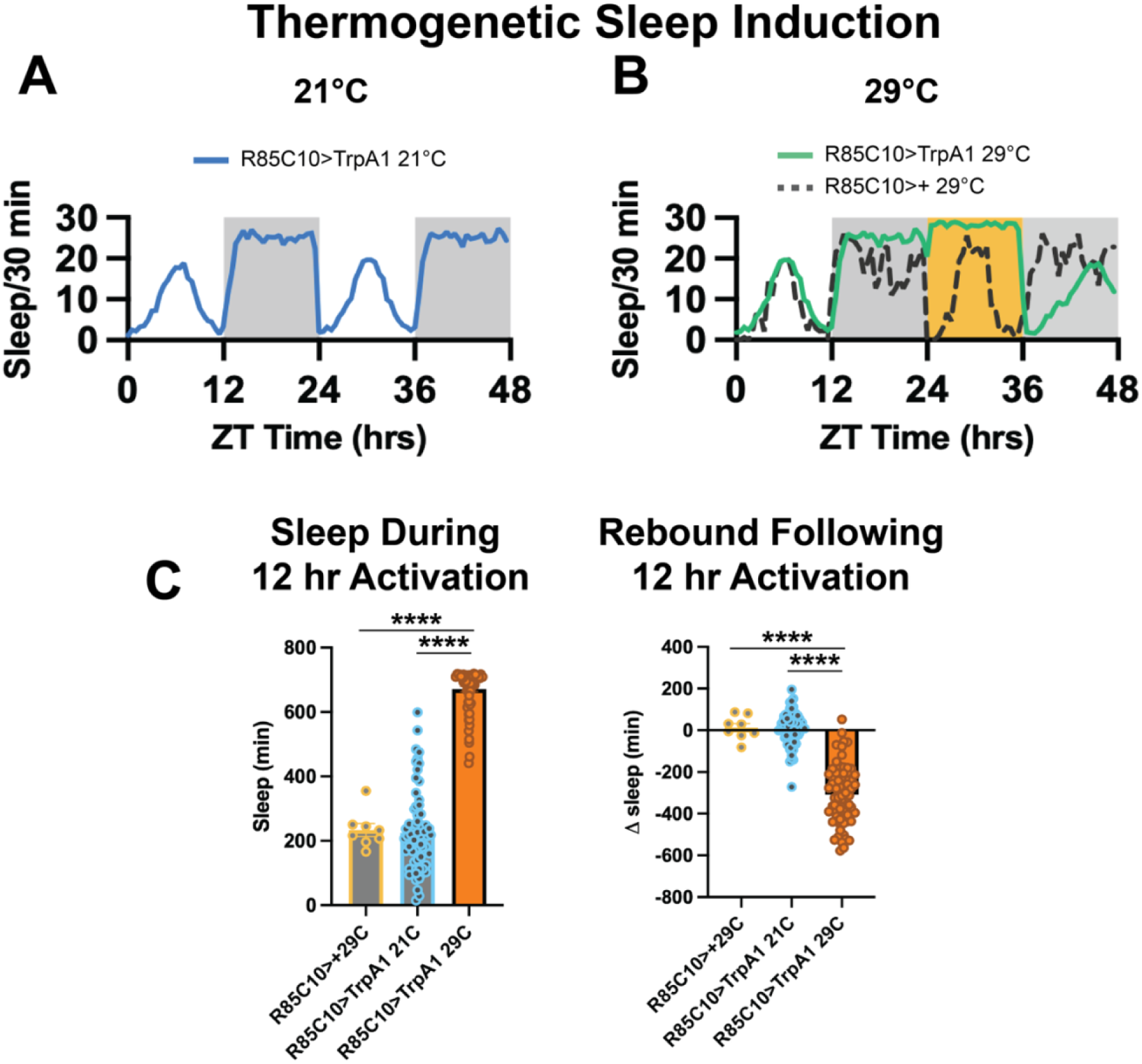
Modulation of sleep using thermogenetic TrpA1 activation. Figures show a GAL4 line (R85C10-GAL4) that promotes sleep and the dissipation of sleep drive following activation. (A) Two circadian cycles of sleep behavior at 21°C depicting flies bearing both the GAL4 and UAS-TrpA1 transgenes. (B) Two circadian cycles of sleep behavior for two genotypes: a GAL4 line paired with a UAS-TrpA1 transgene as well as a GAL4/+ line. The flies are maintained at 21°C for most of the experiment except for a 12 hr transition to 29°C at the beginning of the second day (depicted in orange). (C) Quantification of total sleep during the 12 hr activation for each condition (left) and the change in sleep during the 12 hr rebound period following activation compared to the previous day (right). **** p<0.0001 by ANOVA with Tukey’s multiple comparisons.

**Figure 3.**
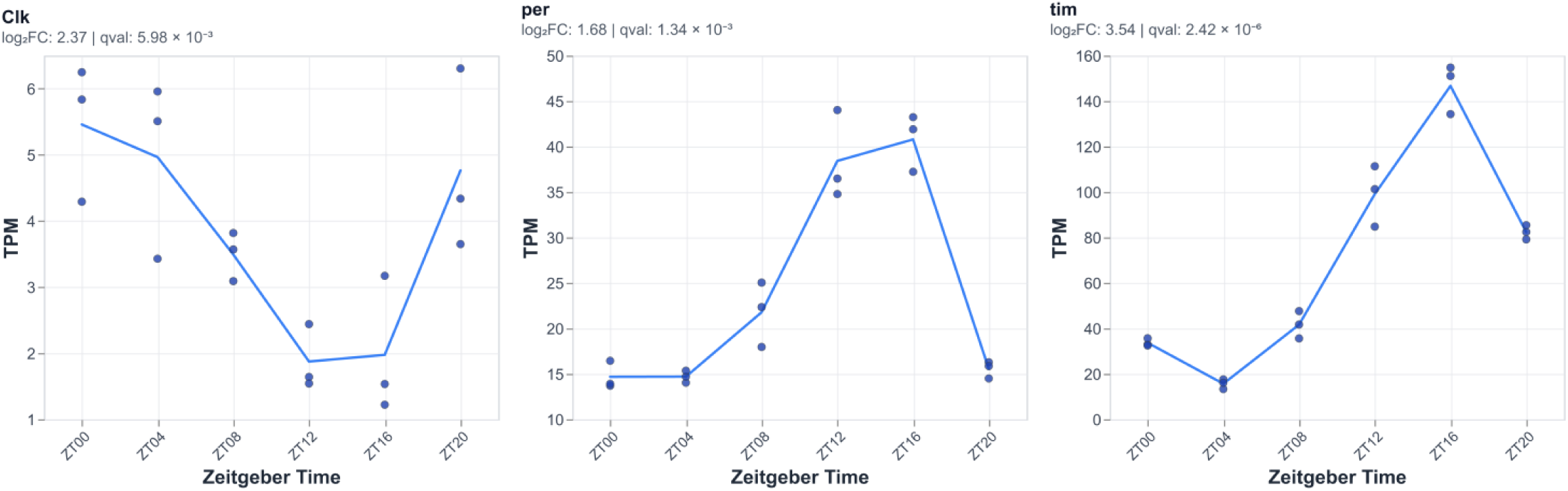
Daily oscillations of core clock-gene expression across Zeitgeber time. Transcript abundance (TPM) is shown for three canonical circadian genes (Clk, per, tim) measured every 4 hours between at ZT00 - ZT20. Points denote individual biological replicates at each timepoint; the connecting line represents the mean TPM across replicates. For each gene, rhythmicity across Zeitgeber time was tested using the RAIN (Rhythmicity Analysis Incorporating Nonparametric methods*) algorithm, and the estimated differential amplitude (log2 FC) and false-discovery-adjusted q-value are reported above each panel.

### Identification of Consistent Differentially Expressed Sleep and Wake Genes Across Datasets

To identify comparable high amplitude, sleep homeostatic genes, we identified differentially expressed genes in each dataset with a log_2_ fold-change threshold of >1, which is still below that observed for whole brain core circadian gene oscillations (Fig. 3). Interestingly, we also note that the number of differentially expressed genes for mechanical SD peaked at 6h of sleep deprivation (for log_2_FC1, 321 genes v. 57 for MechSD3 and 39 for MechSD12). We hypothesize that after 6h of SD, homeostatic set point thresholds may be triggered reversing these transcriptional changes resulting in fewer differentially expressed genes after 12h of SD. This is consistent with predicted homeostatic kinetics^44^. Thus, a cleaner molecular assessment may be best visualized using shorter SD stimuli^45^.

To identify genes that respond to sleep-wake manipulations across datasets, we selected genes that both change in at least 2 datasets and exhibit changes that are consistent with respect to sleep-wake state, i.e., if a transcript rises with wake in one condition, it also rises with wake (or falls with sleep) in a second dataset. Overall, we identify 98 consistent wake genes, i.e., up regulated in wake, and 16 consistent sleep genes, i.e., up regulated in sleep (Table S1). While most differentially expressed genes are non-overlapping between two datasets, we found that the overlap between many different datasets was greater than by chance. This suggests sleep- rather than method-related signal (Fig. 4A). We find that significant overlaps were evident using similar methods of sleep deprivation suggesting that the transcriptomic response may be related in part to the method rather than the sleep deprivation. For example, for wake genes for which there are significant numbers, the pairwise overlaps between the three mechanical sleep deprivation (MechSD3, 6, 12h) conditions ranges from 79-267x more than by chance.

**Figure 4.**
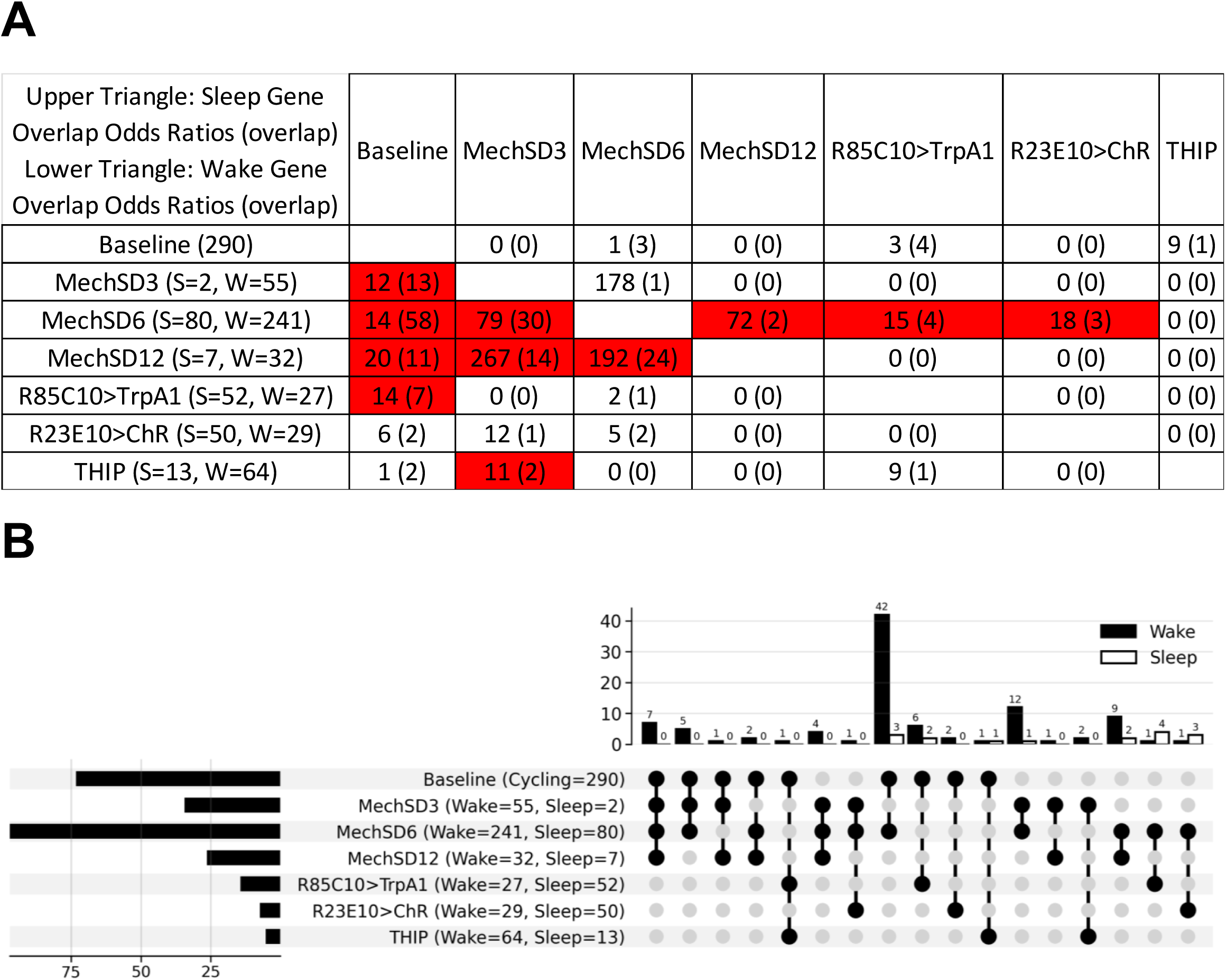
Shared transcriptional signatures of high-amplitude sleep- and wake-regulated genes across experimental paradigms. A. Overlap enrichment of common high-amplitude genes (|log₂FC| > 1, qvalue < 0.05) between pairs of experiments, assessed using one-sided Fisher’s exact test. The matrix is arranged with sleep-gene overlap statistics in the upper triangle and wake-gene overlap statistics in the lower triangle; each cell shows the rounded odds ratio followed by the overlap count in parentheses (e.g., *79 (23)*). Row headers indicate the total number of sleep (S) and wake (W) regulated genes for each experiment, and cells with BH-adjusted qvalue < 0.05 are highlighted in red. B. UpSet plot summarizing intersections of high-amplitude, consistently wake- and sleep-associated transcripts (|log₂FC| > 1.0, qvalue < 0.05) across the seven whole-brain wake and sleep paradigms. Total differentially expressed gene counts per condition are shown horizontally, and intersection sizes are shown vertically. Most high-amplitude wake genes (42) overlapped with baseline cycling transcripts and were upregulated in MechSD6, whereas high-amplitude sleep genes showed a more even distribution of overlaps across paradigms.

Nonetheless, we also observed examples where distinct methods of sleep manipulation exhibited significant overlap suggesting the presence of sleep-related biological signals, i.e., homeostatic genes independent of method. For example, we observe significant overlap (11x) in regulated genes between mechanical sleep deprivation (3h) and THIP treatment for genes up regulated during wake. In addition, we found overlap for sleep up regulated genes between mechanical SD 6h and both thermogenetic (R85C10/TrpA1; 15x) and optogenetic (R23E10/ChR +ATR; 18x) activation (Fig. 4A). Notably, we also find substantial overlap between baseline sleep and both mechanical SD (12-20x), on the one hand, and thermogenetic sleep promotion (R85C10/TrpA1; 14x) on the other, underscoring the notion that homeostatic mechanisms operate under baseline conditions (Fig. 4A). We also examined dataset overlap using UpSet plots, further underscoring the presence of genes that are not only consistent across datasets, including those within the mechanical SD datasets, but also the overlap between mechanical and other sleep-wake manipulations (Fig. 4B). Using these criteria, we find that these consistent sleep-wake genes are at most evident in just 4 datasets (out of 7 examined) indicating that there do not appear to be universal high amplitude transcriptomic markers of sleep homeostasis evident in the whole brain.

Using DAVID, we identified gene ontology categories enriched in these datasets (Fig. 5; Table S2). Among the GO categories enriched among wake genes were mitochondrially related GO terms (e.g., mitochondrial inner membrane) consistent with those observed in the sleep homeostatic neurons, the dorsal fan shaped body (dFB)^12^. In fact, among the 13 such genes, 3 overlap with the 21 mitochondrial inner membrane genes identified from the dFB dataset. This result suggests that the dFB changes are evident in whole brains and may also extend beyond the dFB and is consistent with sleep-wake dependence of oxidative stress more broadly in the brain and even outside the brain^46–48^. We also identified novel pathways including those involved in protein translation and immune/defense response. Immune regulation has been functionally linked to sleep in *Drosophila*^49–51^. We did not identify any significantly enriched GO categories from the sleep up regulated genes.

**Figure 5.**
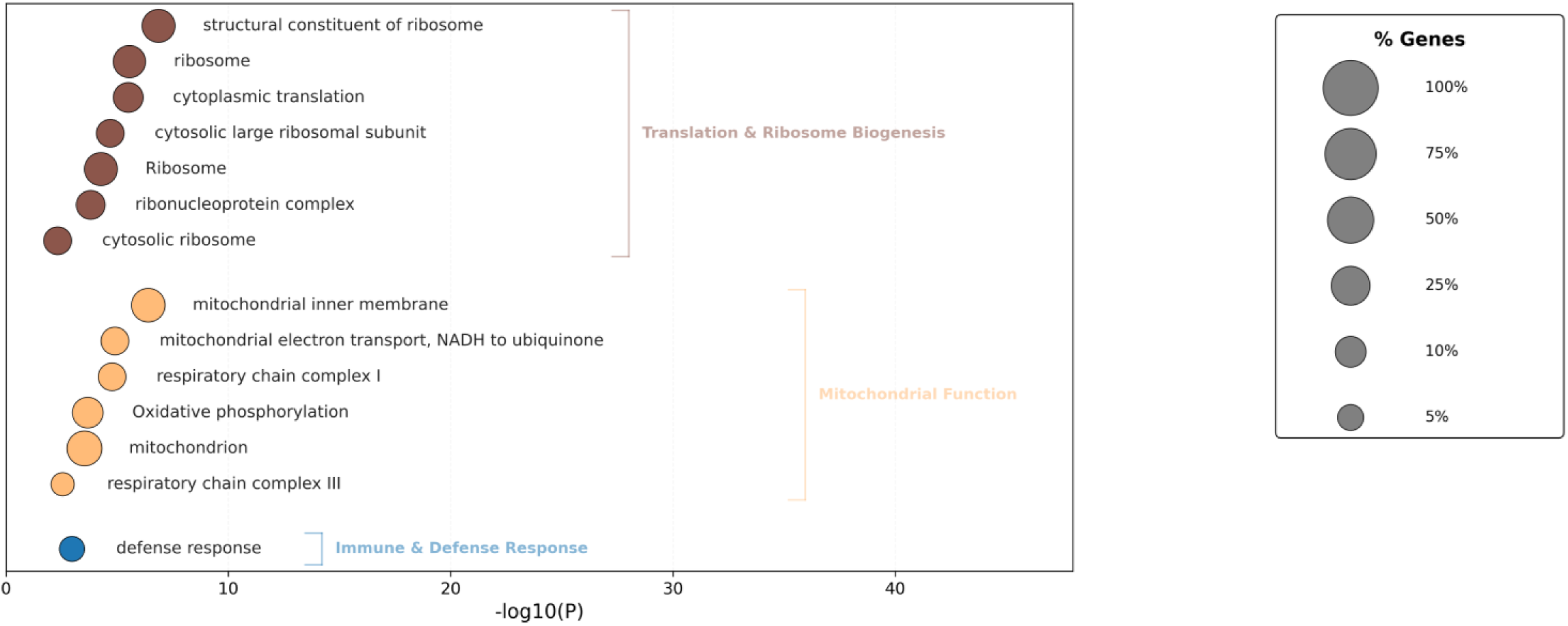
Functional enrichment of high-amplitude consistent wake-up regulated gene sets. Bubble plot showing Gene Ontology (GO) and KEGG pathway enrichment (FDR < 10%) for high-amplitude wake up-regulated transcripts. The x-axis indicates statistical significance (-log10P.) Major functional clusters include Mitochondrial Function (oxidative phosphorylation and electron transport), Translation & Ribosome Biogenesis, RNA Processing, and Immune & Defense Response. Circle sizes represent the percentage of genes enriched within each term.

To determine if these candidate homeostatic markers play an *in vivo* functional role in sleep wake regulation, we developed a GPT based method we term fl.ai (pronounced “fly”), to mine the literature for links between sleep-wake candidate genes and *in vivo* sleep-wake effects (see Methods). This approach aggregates references (Flybase, PubMed, and Europe PMC) and then scrapes full paper text (PubMed Central Open Access, Europe PMC, Unpaywall, OpenAlex, Crossref, and DOI.org Resolver). We then apply GPT5-Nano to generate reference summaries from the text and then GPT5-mini to aggregate the reference summaries and categorize the gene as circadian or sleep related. GPT provides a confidence score based on the level of evidence in support. 21 genes show a circadian association, 8 show a sleep association, and 7 associate with both (see Table S3). For example, the immune-induced secreted Bomanin peptide gene *BomBc2* displays robust up regulation by 3, 6, and 12 h of mechanical sleep deprivation (Table S1). Glial knockdown of *BomBc2* significantly reduces nighttime sleep consistent with an in vivo sleep function^32^, suggesting that wake associated increases in *BomBc2* observed here may subsequently promote sleep, forming a functional homeostatic feedback loop.

As we did not identify any sleep-wake genes that are evident across all datasets, we opted to lower our fold-change threshold to log_2_(0.5) but maintain our rigorous statistical threshold (q<0.05) to see if such genes became evident. Following the pipeline, we identified 647 consistent wake and 141 consistent sleep genes (Table S1). Consistent with the idea that we are capturing more signal versus noise at the lower threshold, we observe many more datasets exhibiting significant overlap between consistent sleep/wake genes (Fig. 6A). We now observe uniform overlap among all three mechanical SD datasets for both sleep and wake genes (36x-286x more than by chance). In addition, we again observe overlap between baseline and mechanical SD datasets (4-7x for wake genes) consistent with shared transcriptomic responses and molecular mechanisms. Most remarkably, we now observe significant and uniform overlap of wake associated genes between baseline (2-7x), ChR (3-8x) and THIP (2-10x) datasets and all of the other datasets, highlighting potential responses independent of the method of sleep-wake manipulation. UpSet plots now show that one gene encoding the ribosomal protein subunit *RpL23* is evident in 6 out of the 7 datasets (Fig. 6C). Wake GO analysis (Fig. 7A; Table S2) revealed a much larger list of enriched categories but continued to include KEGG pathways for Ribosome and Oxidative Phosphorylation as in the log_2_FC1 dataset, including several genes encompassing the Mitochondrial Inner Membrane GO category (Fig. 7B). Among the sleep genes (Fig. 7C), plasma membrane (cellular compartment), ATP binding (molecular function) and calcium ion binding (molecular function) are among the enriched pathways.

**Figure 6.**
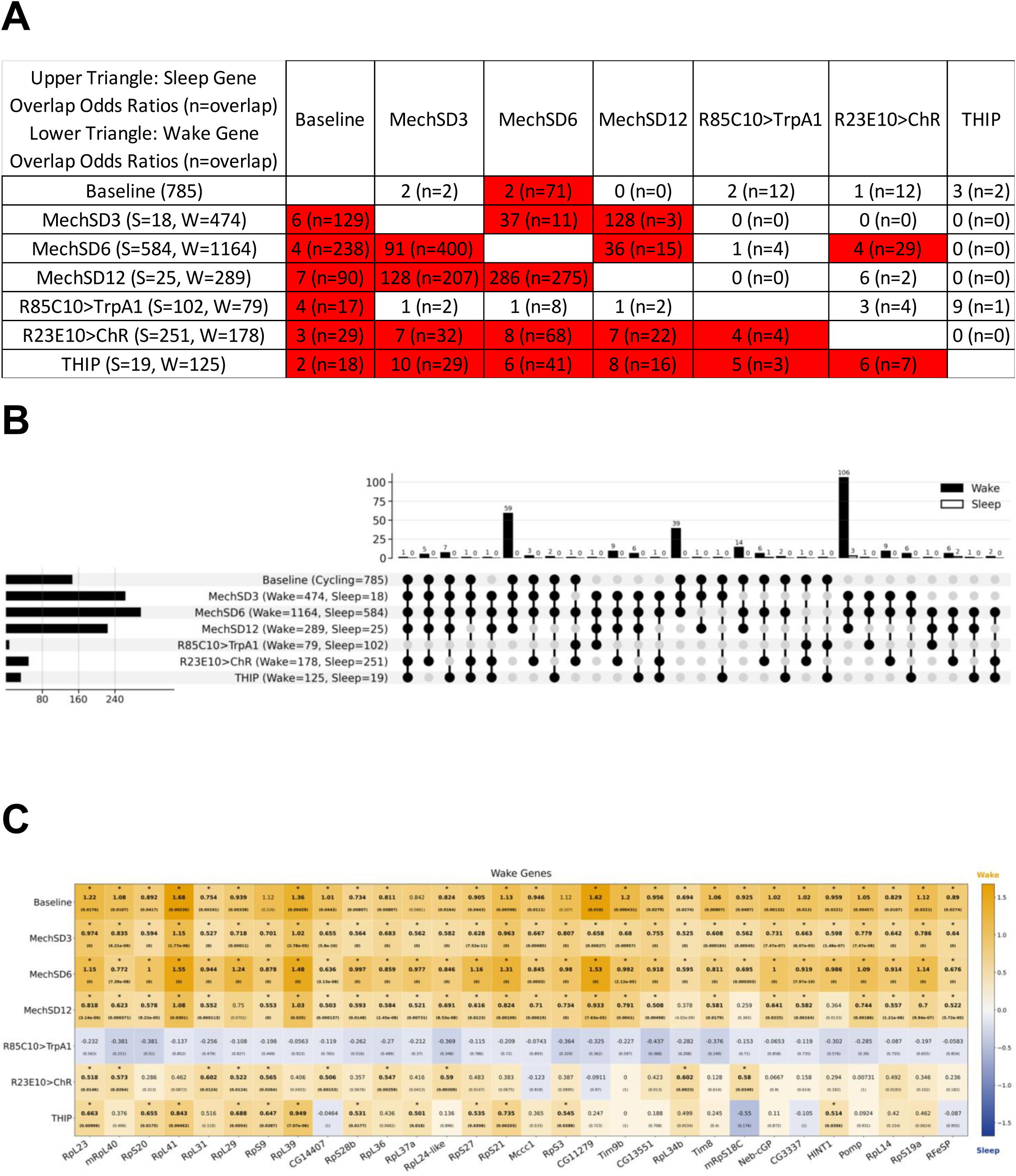
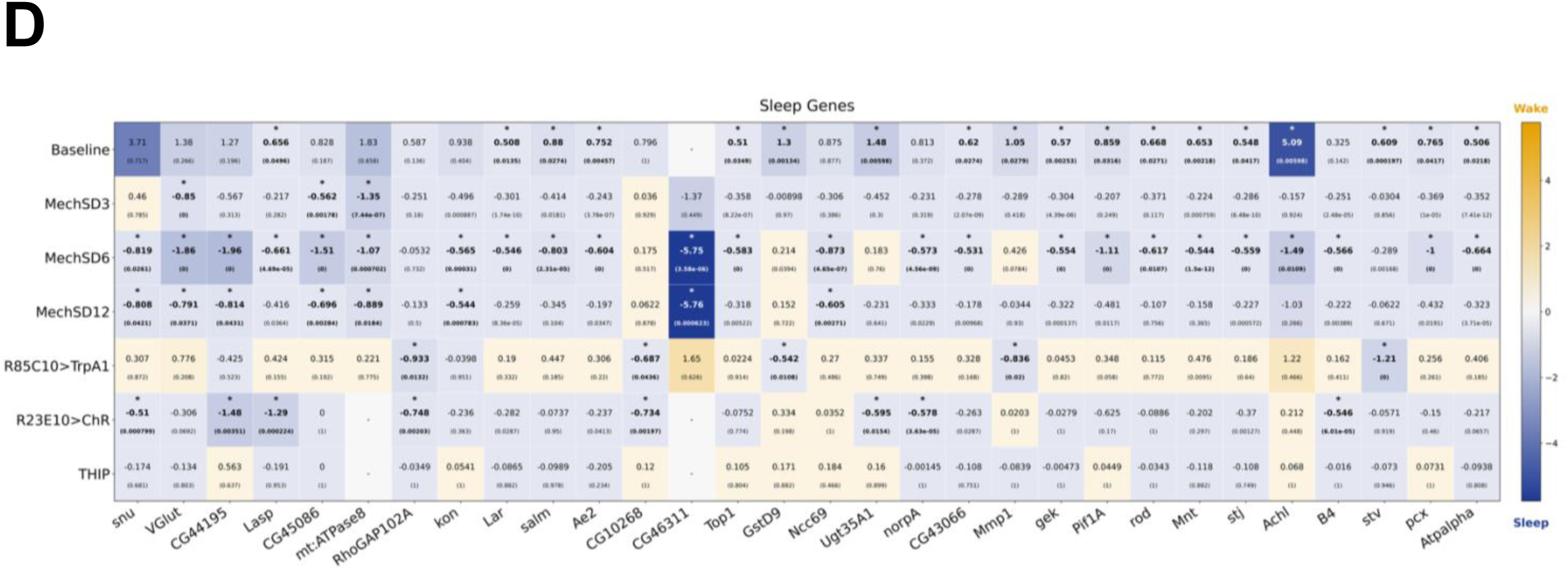
Shared transcriptional signatures of low-amplitude sleep- and wake-regulated genes across experimental paradigms. A. Overlap enrichment of common low-amplitude genes (|log₂FC| > 0.5, qvalue < 0.05) between pairs of experiments, assessed using one-sided Fisher’s exact test. The matrix is arranged with sleep-gene overlap statistics in the upper triangle and wake-gene overlap statistics in the lower triangle; each cell shows the rounded odds ratio followed by the overlap count in parentheses (e.g., *8 (n=16)*). Row headers indicate the total number of sleep (S) and wake (W) regulated genes for each experiment, and cells with BH-adjusted qvalue < 0.05 are highlighted in red. B. UpSet plots summarizing intersections of low-amplitude, consistently wake- and sleep-associated transcripts (|log₂FC| > 0.5, qvalue < 0.05) across the seven whole-brain wake and sleep paradigms, including transcripts present in at least three experiments. Total differentially expressed gene counts per condition are shown horizontally, and intersection sizes are shown vertically. The overlap distribution across sleep paradigms resembled that observed for the high-amplitude sleep- and wake-conforming transcripts, with many wake genes driven by overlap with cycling genes in baseline conditions that were upregulated during wake and mechanical sleep deprivation. C. Heatmap of low-amplitude, consistently wake-associated transcripts (same gene set as in panel B; |log₂FC| > 0.5, qvalue < 0.05). Rows represent the seven whole-brain sleep perturbing paradigms and columns represent wake-associated genes (ranked by recurrence across paradigms); color intensity encodes wake-normalized log₂FC (orange, wake direction; blue, sleep direction; white, near zero). Each cell is annotated with log₂FC (top) and qvalue (bottom), and asterisks indicate experiment-level significance at the stated thresholds. D. Heatmap of low-amplitude, consistently sleep-associated transcripts (same gene set as in panel B; |log₂FC| > 0.5, qvalue < 0.05). Rows represent the seven whole-brain paradigms and columns represent sleep-associated genes (ranked by recurrence across paradigms); color intensity encodes wake-normalized log₂FC (orange, wake direction; blue, sleep direction; white, near zero). Each cell is annotated with log₂FC (top) and qvalue (bottom), and asterisks indicate experiment-level significance at the stated thresholds.

**Figure 7.**
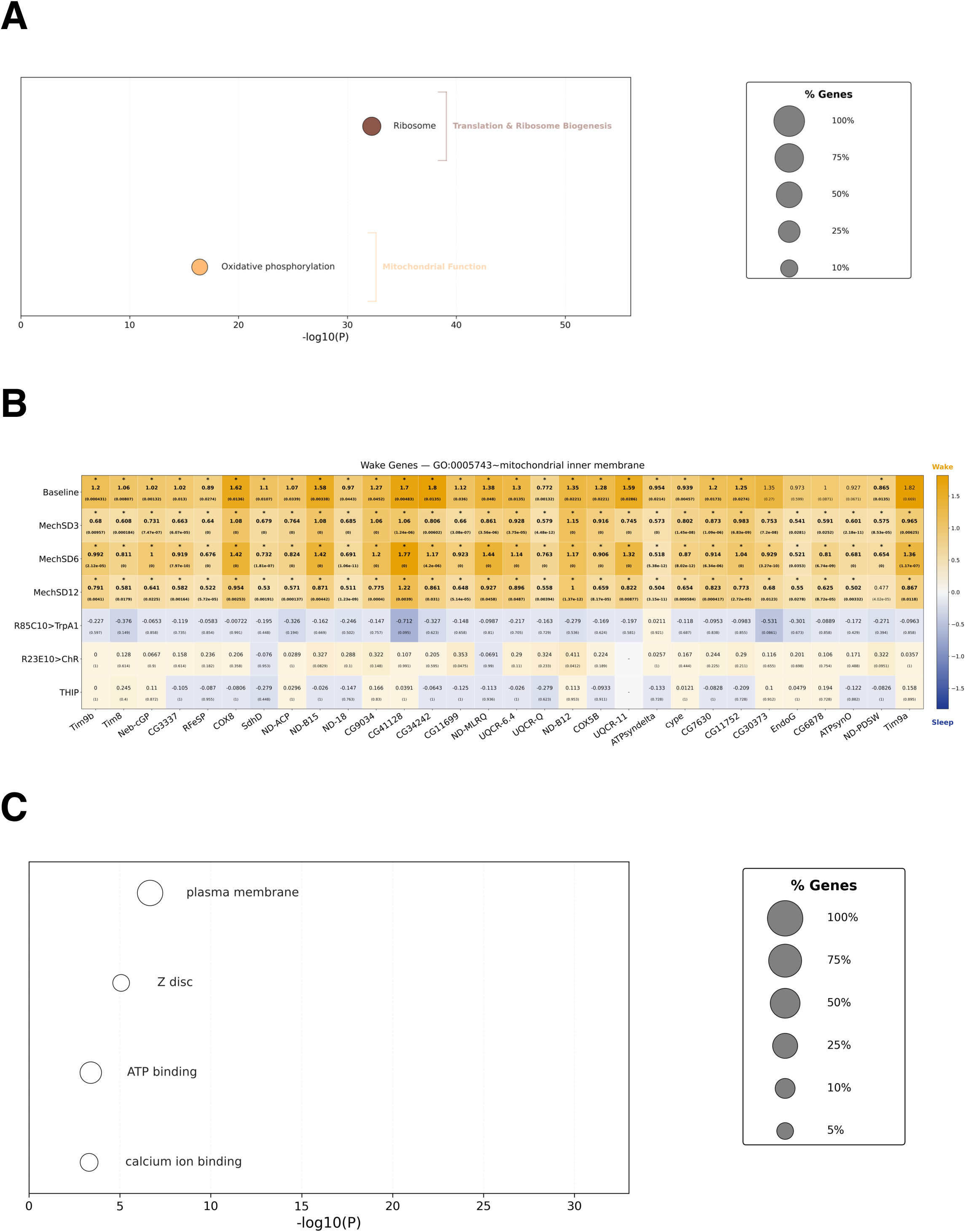
Functional enrichment of low-amplitude consistent wake- and sleep-up regulated and gene sets. A. Bubble plot showing KEGG pathway enrichment (FDR < 10%) for low-amplitude wake up-regulated transcripts. The x-axis indicates statistical significance (-log10P.) Circle sizes represent the percentage of genes enriched within each term. B. Heatmap of low-amplitude wake up-regulated transcripts enriched in the GO Cellular Component term “mitochondrial inner membrane” (GO:0005743; FDR < 0.1 using the DAVID bioinformatics web service). Genes are ordered by the number of datasets in which they reached statistical significance; cell values show wake-normalized log₂FC with q-values in parentheses. Cells are bolded and marked with an asterisk where |log₂FC| > 0.5 and q < 0.05. C. Bubble plot showing Gene Ontology (GO) and KEGG pathway enrichment (FDR < 10%) for low-amplitude sleep up-regulated transcripts. The x-axis indicates statistical significance (-log10P.)

To identify potential upstream regulators that may orchestrate sleep-wake dependent gene regulation, we used the HOMER algorithm^52^ to identify sequences enriched in genomic regions up to 1kb upstream of their transcription start sites (Fig. S1). Among the most significant hits among wake induced genes includes the palindromic E box sequence CAGCTG, a classical binding site for basic helix-loop-helix transcription factors. We identified four such factors among our wake-induced genes including *bigmax*, *HLH3B*, *E(spl)m3-HLH*, and *E(spl)mbeta-HLH* (Table S1), suggesting these factors may be direct upstream regulators of wake-induced gene expression.

When we ran *fl.ai*, we found that 170 of the wake genes had a circadian (110), sleep (39) or both (21) association and 46 of the sleep genes (23 circadian, 7 sleep, 16 both). Notable are neuropeptide genes such as Pigment Dispersing Factor (PDF) and SIFamide (SIFa) as well as a host of mitochondrially encoded genes. PDF and SIFa were both induced after 3 and 6h of mechanical sleep deprivation. In addition to its role in coordinating circadian rhythms, PDF promotes wakefulness^53–55^, suggesting the transcriptional response further sustains wakefulness. On the other hand, SIFa promotes sleep^56, 57^ suggesting that the transcriptional response supports homeostasis, i.e., wake-induced up regulation of *SIFa* transcript in turn will drive sleep. Several other neuropeptide genes are also wake-induced and neuropeptide hormone activity is an enriched wake pathway (Table S2). Notably, among the sleep up regulated genes are three members of the NARROW ABDOMEN (NA) cation leak channel complex including *na* itself as well as regulatory subunits *unc79* and *unc80*, each of which exhibit time-dependent oscillation peaking at the end of the sleep period while both are down regulated in MechSD6 (Table S1). As is the case for PDF, the NA/UNC79/UNC80 channel complex is important for circadian rhythms of neuronal activity in clock neurons as well as circadian behavior^58–60^. In addition, *unc79* knockdown results in elevated sleep, even under wake-inducing starvation conditions^61^. Thus, the coordinated expression of sleep induced expression of *unc79*, as well as *na* and *unc80*, may result in subsequent wakefulness, consistent with a homeostatic feedback role. In contrast, *vGlut* is induced by sleep, yet may play a non-homeostatic role in sleep maintenance. Knockdown of *vGlut* in a subset of sleep-promoting dFB neurons significantly reduces sleep^62^. Taken together, these findings suggest a complex interplay between homeostatic forces and state-sustaining forces. Moreover, our analysis has identified several transcript markers of sleep-wake history consistent across at least 2 datasets which have been validated with an in vivo sleep-wake role consistent with a homeostatic function.

### Identification of Transcripts Correlated with Sleep-Wake History and/or Rebound

Given that our preliminary analysis did not identify a candidate sleep-wake or homeostatic marker across all datasets, we complemented this approach to see if there were markers that correlate with sleep-wake and/or sleep rebound integrating data across all of our datasets. Here we also include four TrpA1 lines whose controls showed acute temperature dependent effects on sleep and TrpA1 dependent sleep rebound effects. Previous studies have demonstrated that wakefulness can be thermogenetically induced via activation of TrpA1; however, not all wake-promoting driver lines can induce a subsequent rebound^63^. We identified four lines, collectively referred to as WakeGAL4s, that showed significant wake promotion after 12 h activation (Fig. S2). Here we used briefer 3h activation to identify acute activation effects and observed significant rebound with HC-GAL4^64^, but not with three other lines (R21H11-GAL4, R60F09-GAL4, R65H02-GAL4; Fig. S3A-D). A closer analysis of the HC-GAL4 line suggests that it drives expression in previously described wake-promoting *pickpocket* peripheral neurons^65^ and wake and rebound effects observed during thermogenetic activation of HC-GAL4 neurons could be blocked by GAL80 expression in *pickpocket* neurons (Fig. S4). As we observed acute thermal effects even in non-TrpA1 controls, we could not observe significant wake effects in TrpA1-expressing flies during this briefer activation relative to 29°C controls (Fig. S4). As a result, we did not include these datasets in our initial differential gene expression analysis but did include them in our correlation analyses here. We also removed two published datasets, optogenetic and THIP^41^, for which we did not have direct sleep data to perform correlations. We then asked if there were genes whose expression correlated with sleep-wake history prior to sampling and/or sleep rebound after sampling across these 9 datasets. We also considered the possibility that gene expression may integrate sleep-wake experience over different time scales. Thus, we also considered correlations over distinct bins of sleep wake history from 0-1h prior to 0-12h prior or sleep rebound 0-1h after to 0-12h after. We used Pearson correlation to identify linear correlations between sleep/sleep rebound amount and log2 normalized gene expression (transcripts per million reads). We also corrected for false discovery and selected genes using a stringent q<0.05 threshold.

Using this approach, we identified 149 genes that positively correlate with sleep history (i.e. genes that may be up regulated by sleep) and 66 genes that negatively correlate with sleep history (i.e., genes that may be up regulated by wake: Table S4). To better visualize the data, we created an expression heat map and clustered genes and samples by similarity (Fig. 8). Using this approach, we find that samples show clear clustering based on temperature with 21°C, 25°C, and 29°C samples clustering together irrespective of sleep-wake condition. Nonetheless, we observe clustering of mechanical sleep deprivation samples whether 3, 6 or 12h taken at ZT0 separately from baseline ZT0 samples. Within the 29°C cluster, we observe separation of GAL4/TrpA1 from GAL4/+ consistent with activation-dependent gene expression. Notably, the sleep promoting R85C10/TrpA1 at 29°C clearly separates from all other 29°C samples including other wake GAL4/TrpA1 lines, indicating a transcriptomic response in this sleep promoting line distinct from controls and other 29°C samples (both GAL4/TrpA1 and GAL4/+). Interestingly, R85C10/TrpA1 clusters with other 21°C samples, suggesting that activation may mimic the sleep-inducing effects of cooler temperatures^66, 67^ at the transcriptome level. Thus, the correlation approach appears to identify sleep-wake relevant transcriptomic responses.

**Figure 8.**
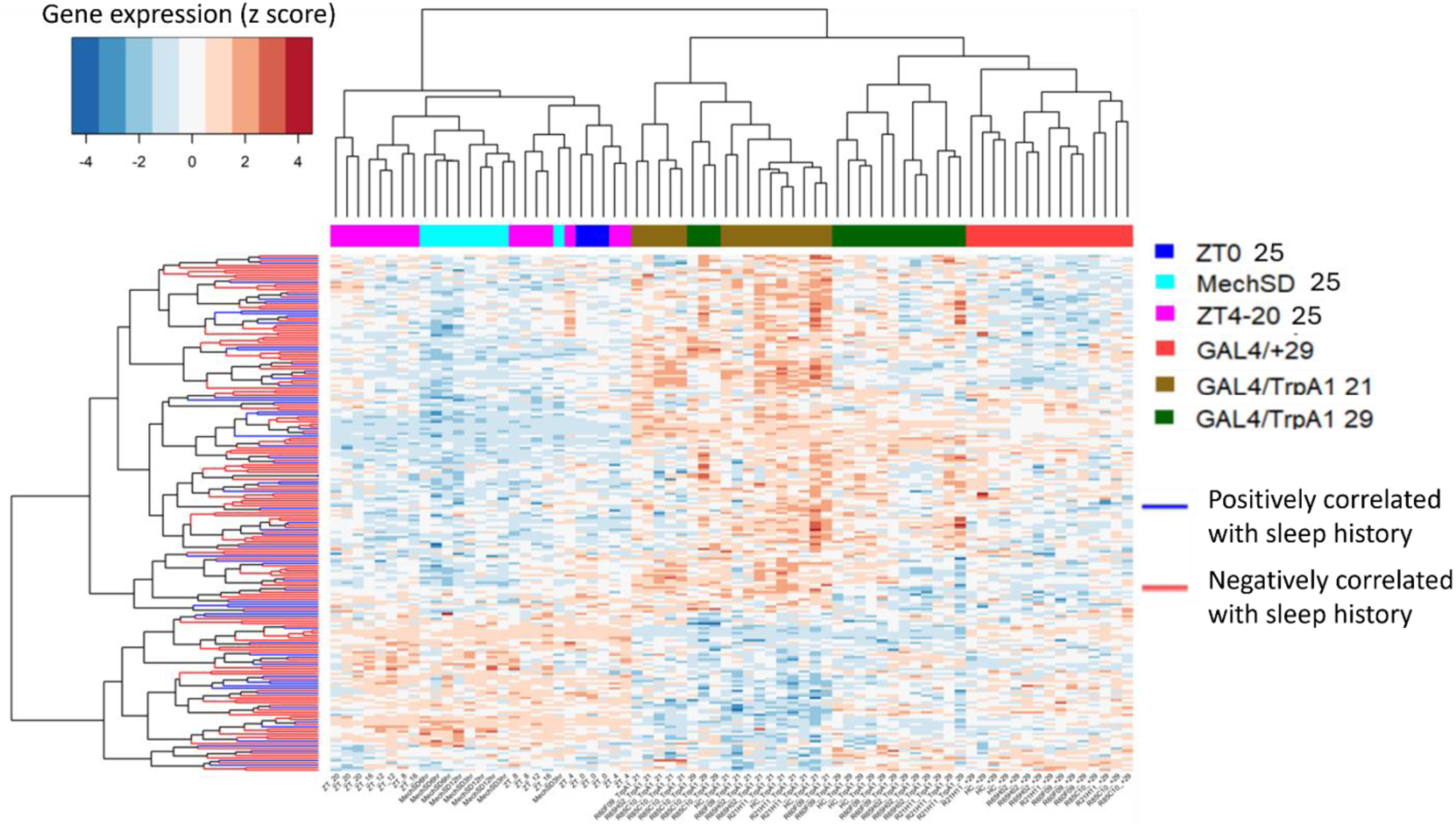
Gene and Sample Clusters Based on Sleep–Wake History. Each row represents a gene, and each column represents a sample. Gene expression values were z-score transformed across samples to visualize relative up- and down-regulation. Rows (genes) and columns (samples) were independently clustered using hierarchical clustering based on Pearson correlation. Gene–gene and sample–sample similarity matrices were calculated as *(1 – Pearson correlation)*, converted to distance matrices, and clustered using the complete linkage method. The resulting dendrograms illustrate patterns of co-expression among genes and similarity relationships among samples. Heatmap clustering of genes and samples reveals a strong effect of temperature, with samples at 25 °C clustering on the left, 21 °C in the middle, and 29 °C on the right. From a sleep–wake perspective, mechanical sleep deprivation (MechSD) clusters separately from ZT0 samples. In contrast to the PCA analysis, sleep-promoting R85/TrpA1 samples cluster separately from other wake TrpA1 samples and instead group with 21 °C samples. Additionally, wake TrpA1 samples at 29 °C cluster apart from wake control samples at the same temperature, despite similar sleep–wake states, suggesting genotype- or activity-dependent effects beyond sleep state alone.

To identify genes that are similarly regulated by cooler ambient temperature (21°C) and by R85C10 neuronal activation (29°C), we first defined putative thermally regulated genes as those differentially expressed between WakeGAL4/+ 29°C control and WakeGAL4>TrpA1 21°C (Table S5). In addition, we examined R85C10/TrpA1 29°C versus R85C10/+ 29°C to examine R85C10 activity dependent gene expression. We then selected overlapping genes between those two sets as putative candidates for R85C10 activation gene regulation that mimic cooler temperature. We then overlaid this list with our sleep correlated gene list. Notably, we find two neuropeptide receptor genes *AstA-R2* and *AstC-R1* that correlate with sleep history and/or rebound and are either up or down regulated under cooler temperatures and R85C10 activation (Table S5). Allatostatin-A and -C (AstA and AstC) are neuropeptides that play important roles in both thermal and sleep responses in part as key neuropeptides for the thermosensory lateral posterior neurons (LPNs) ^68–72^ and circadian clock dorsal neurons 3 (DN3s^73^).

Plotting the correlations across time bins from 0-1 to 0-12h for both positively and negatively correlated genes, we find gene subsets which display stronger correlations over shorter bins while others display stronger correlations over longer bins (Fig. 9). We hypothesize that the expression of the former integrates sleep-wake experience over shorter time scales while the latter integrates over longer time scales. Gene ontology analyses of genes upregulated after wake (Table S6) showed genes involved in immune defense (Fig. 10A). Genes upregulated after sleep (Fig. 10B) counterintuitively revealed a dominant role for mitochondrial pathways, including those associated with the mitochondrial inner membrane. Our prior analysis (Fig. 7B) and published work^12^ suggested that these genes linked to the mitochondrial inner membrane are up regulated after wake, while this correlation analysis shows they are up regulated after sleep. To resolve this potential discrepancy, we compared the gene lists and found that they were entirely non-overlapping suggesting sleep and wake target the mitochondrial inner membrane in discrete ways. Another enriched GO term is solute:sodium transporters (Fig. 10) consisting of the SLC5A transporters *rumpel*, *bumpel*, and *kumpel*. These transporters are expressed in ensheathing glia and are thought to transport glucose to neurons to drive ATP production^74^. Notably, mutants of *rumpel* display reduced daytime sleep^74^. Thus, wake dependent increases in these transporters may function to promote sleep closing a homeostatic loop.

**Figure 9.**
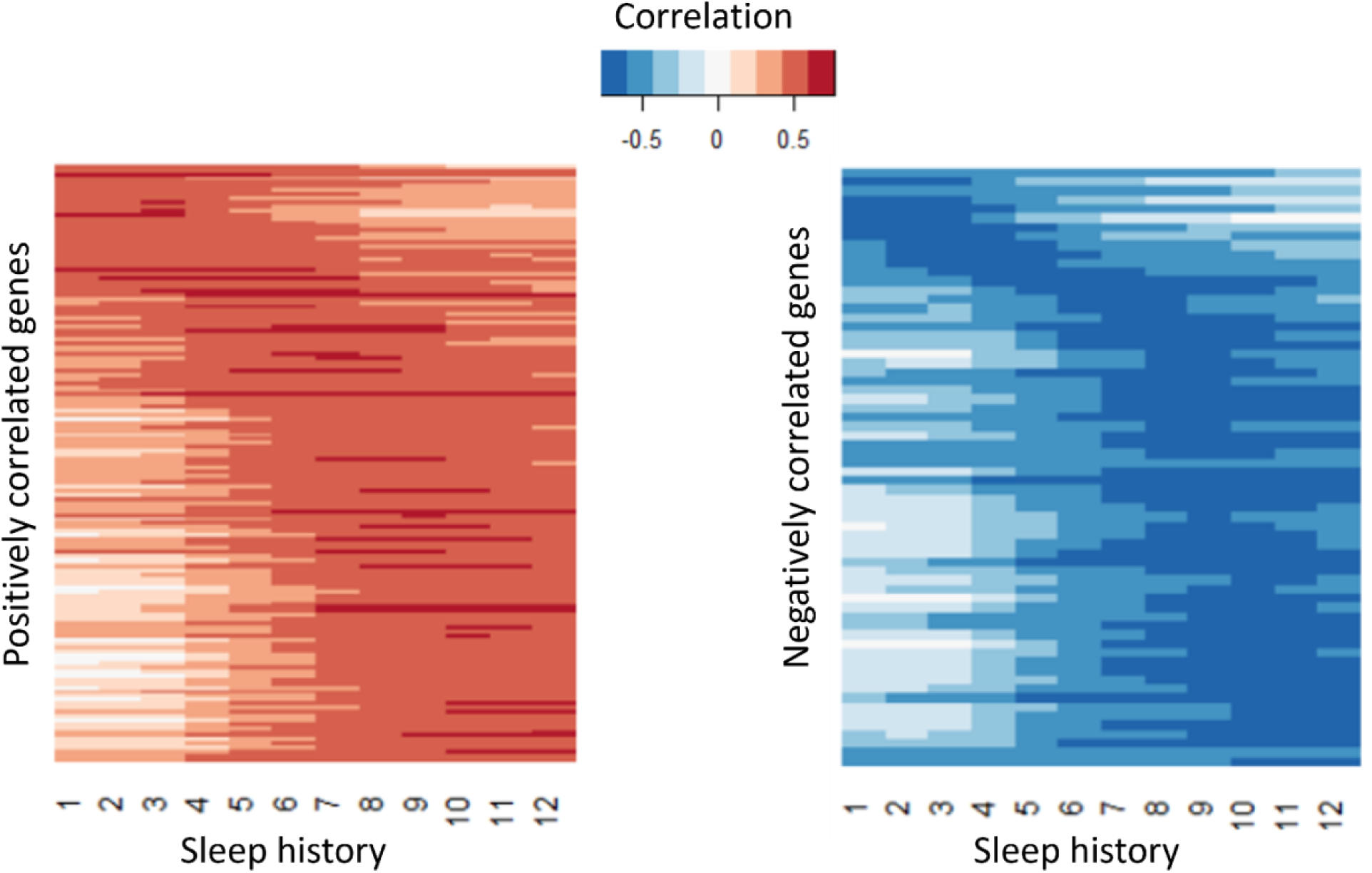
Genes positively or negatively correlated with recent or long-term sleep history. Pearson correlations were calculated between gene expression levels and amount of sleep prior to sample collection. Correlations were evaluated across time bins ranging from 0–1 h to 0–12 h before sampling. Gene expression values (log₂ TPM) were averaged across three biological replicates, and sleep measurements were averaged across the number of flies per sample. A total of 149 genes showed significant positive correlation (correlation coefficient > 0, BH-adjusted *q* < 0.05), while 66 genes showed significant negative correlation (coefficient < 0, BH-adjusted *q* < 0.05). Among both positive and negatively correlated genes, some exhibited stronger associations with shorter sleep-history bins, whereas others showed stronger correlations with longer time windows.

**Figure 10.**
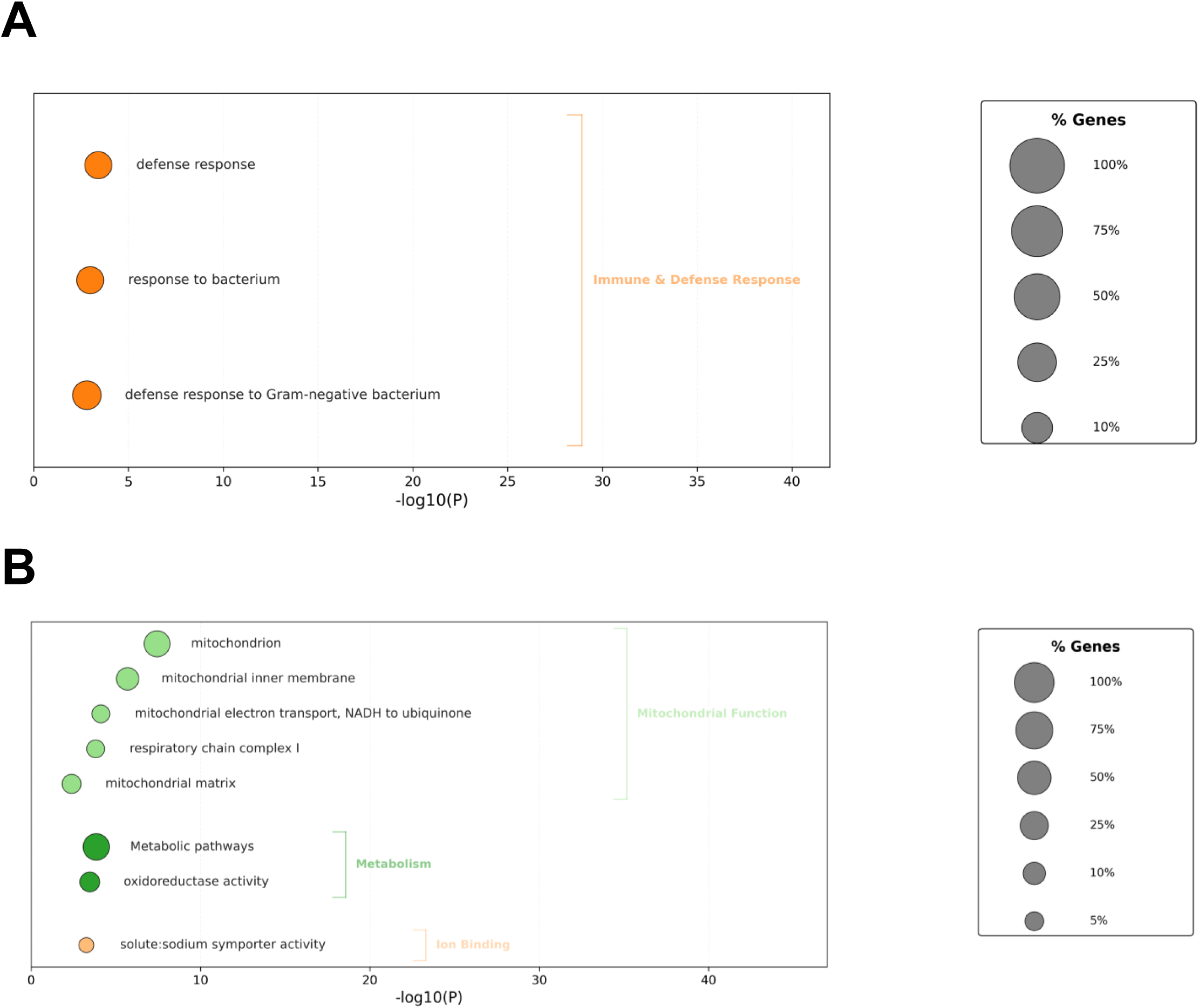
Functional enrichment of transcripts negatively- and positively-correlated with sleep history. A. Bubble plot showing biological process enrichment (FDR < 10%) for genes negatively correlated with sleep-history (Pearson correlation, qvalue < 0.05). The x-axis indicates statistical significance (-log10P.) Circle sizes represent the percentage of genes enriched within each term. B. Bubble plot showing Gene Ontology (GO) and KEGG pathway enrichment (FDR < 10%) for genes positively correlated with sleep-history (Pearson correlation, qvalue < 0.05). The x-axis indicates statistical significance (-log10P.) Circle sizes represent the percentage of genes enriched within each term.

We also performed a parallel analysis for gene expression that correlates with sleep rebound, i.e., the difference in sleep amount before and after a perturbation. Here we identify 63 genes that demonstrate a positive correlation (more sleep rebound, more gene expression) and 258 genes that exhibit a negative correlation (less sleep rebound, more gene expression; Fig. 11, Table S4). As for sleep history we also observe genes that exhibit correlations over shorter term rebound and others over longer term rebound periods (Fig. 12). GO analysis (Table S6) reveals a role for immune-related defense pathways that are positively correlated with wake after sampling (Fig. 13). Given that immunity is also positively correlated with wake prior to sampling, it suggests a general role of immune genes in wake regulation rather than a homeostatic function. We then examined the overlap between genes correlated with sleep history and also sleep rebound (Table S4). We identified 15 genes that are positively correlated with sleep prior to sampling but negatively correlated with sleep after sampling suggesting that these genes reflect sleep-wake experience and can induce a homeostatic response. 5 additional genes are negatively correlated with sleep history and positively correlated with rebound. Remarkably there were no genes that exhibited a positive correlation both before and after sampling or a negative correlation before and after suggesting that these associations reflect true biological signals. One of these genes, *AstA-R2*, has an established role in sleep regulation^75^ (Fig. 14, Table S7). *AstA-R2* expression has a positive correlation with sleep history and a negative correlation with sleep rebound, suggesting that *AstA-R2* declines with wake and this may then trigger subsequent sleep rebound. Knockdown of *AstA-R2* in blood-brain barrier glia elevates daytime sleep, consistent with a wake promoting role^75^. *Trh*, the rate-limiting enzyme for serotonin biosynthesis, is similarly positively associated with sleep–wake history and negatively associated with sleep rebound. Notably, loss of *Trh* leads to both reduced baseline sleep and diminished rebound sleep^76^. In parallel, serotonin levels track with sleep pressure^77^, supporting a model in which *Trh* regulation drives wake history–dependent serotonin production, which in turn promotes rebound sleep.

**Figure 11.**
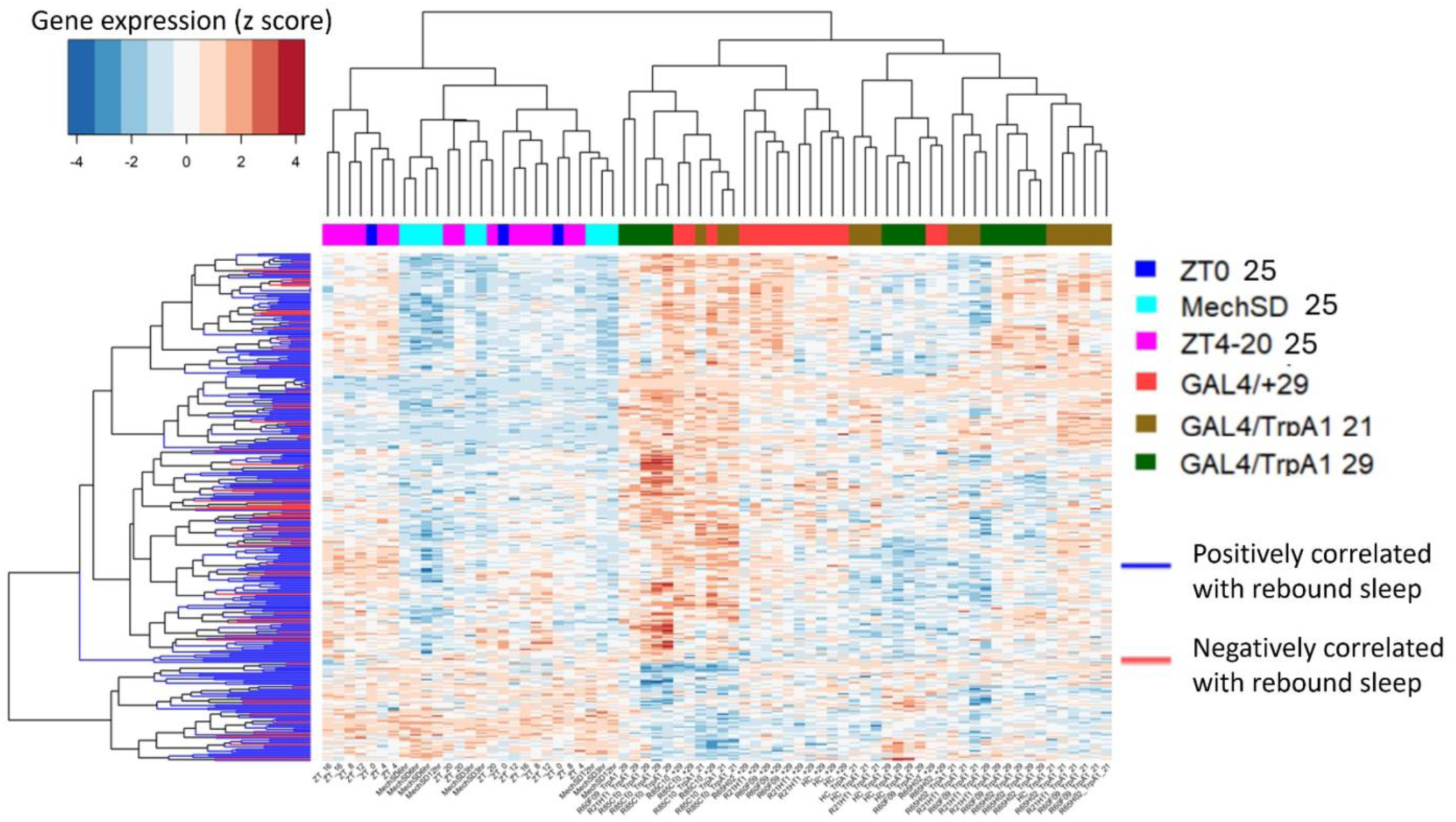
Gene and sample clustering based on sleep rebound–associated genes. Each row represents a gene positively or negatively correlated with sleep rebound, and each column represents an individual sample. Gene expression values (TPM) were z-score normalized across samples to visualize relative up- and down-regulation patterns. Rows (genes) and columns (samples) were independently clustered using hierarchical clustering based on Pearson correlation. Gene–gene and sample–sample similarity matrices were computed as (*1 − Pearson correlation*), converted to distance matrices, and clustered using the complete linkage method. The resulting dendrograms reveal patterns of gene co-expression and similarity relationships among samples. In contrast to sleep history–associated genes, samples collected at 29 °C and 21 °C show greater intermixing, indicating weaker separation by temperature condition.

**Figure 12.**
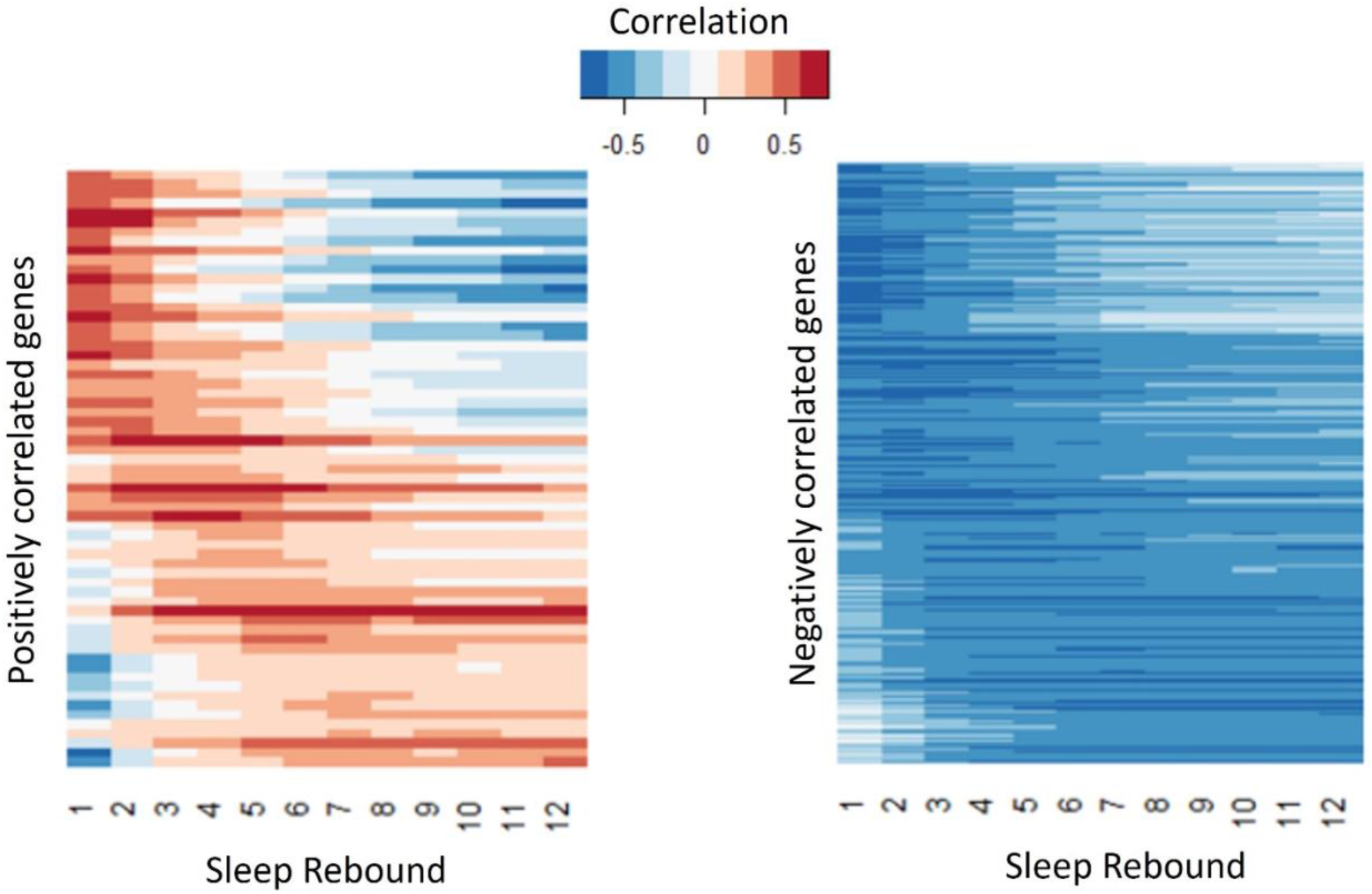
Genes positively or negatively correlated with acute or longer-term sleep rebound. Sleep rebound was defined as the difference between cumulative sleep measured after sampling (1–12 h) and the corresponding cumulative baseline sleep over matched time intervals (1–12 h) for each condition. Pearson correlations were calculated between gene expression levels and each of the 12 rebound intervals. Circadian-control samples were excluded from this analysis. Gene expression values (log₂ TPM) were averaged across three biological replicates, and rebound sleep measurements were averaged across the number of flies per sample. A total of 63 genes showed significant positive correlations (correlation coefficient > 0, BH-adjusted *q* < 0.05), whereas 258 genes showed significant negative correlations (correlation coefficient < 0, BH-adjusted *q* < 0.05). Similar to sleep history correlations, some genes exhibited stronger associations with shorter rebound intervals, while others showed stronger correlations across longer rebound periods.

**Figure 13.**
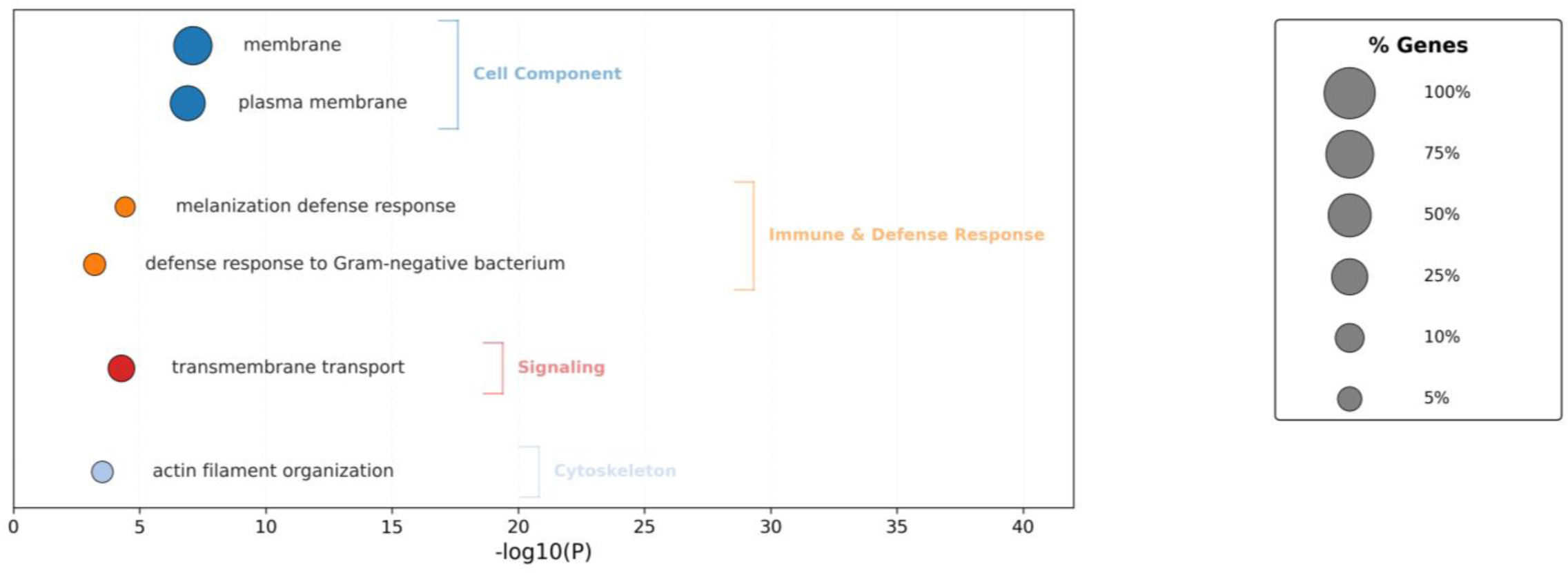
Functional enrichment of transcripts negatively-correlated with sleep rebound. Bubble plot showing Gene Ontology (GO) and KEGG pathway enrichment (FDR < 10%) for genes negatively correlated with sleep-rebound (Pearson Correlation, qvalue.< 0.05). The x-axis indicates statistical significance (-log10P.) Circle sizes represent the percentage of genes enriched within each term.

**Figure 14.**
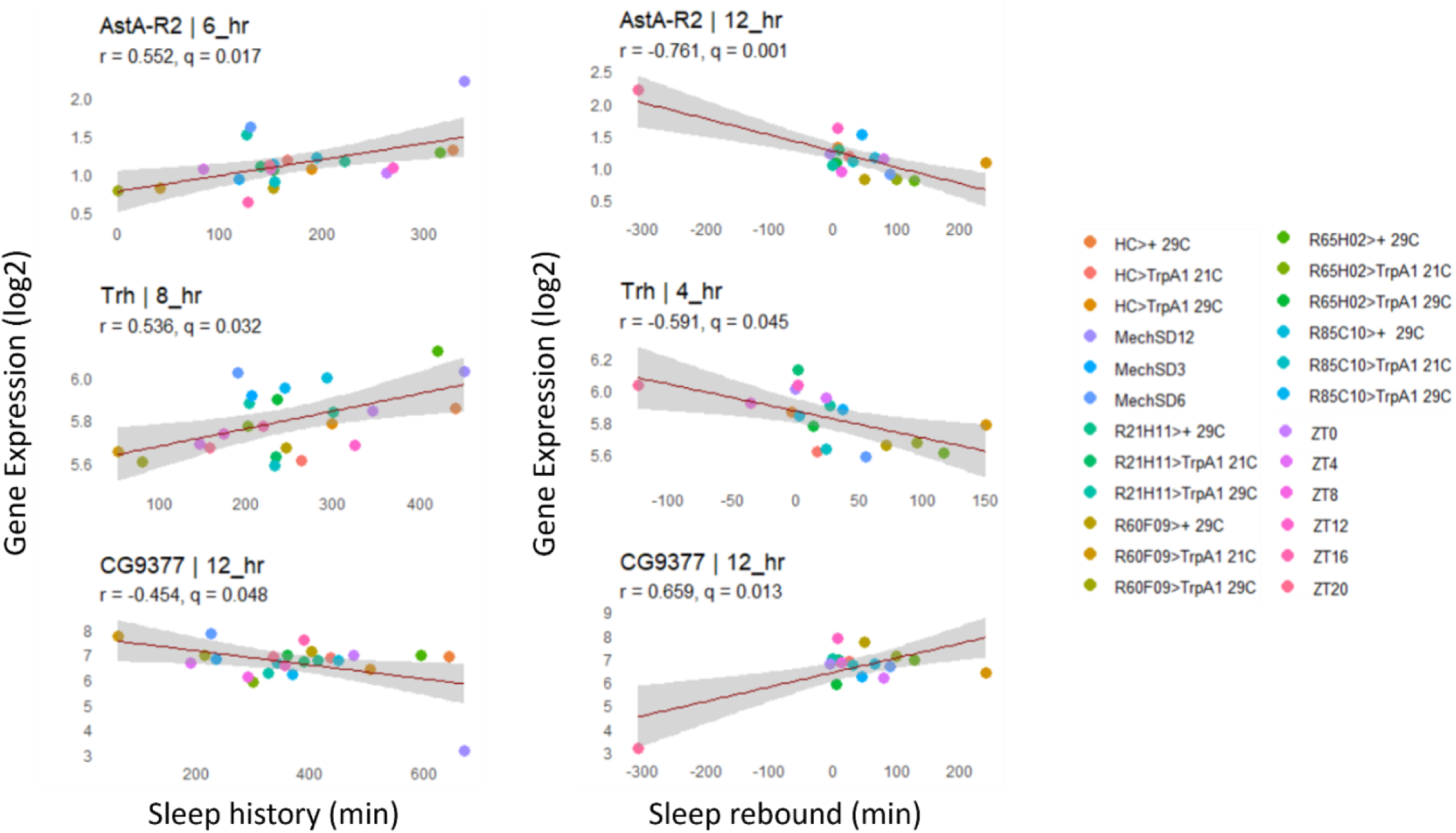
Sleep vs. gene expression plot: Relationship between sleep history and rebound sleep for each genotype plotted against corresponding gene expression levels. In the examples shown, AstA-R2 and Trh expression exhibits a positive correlation with sleep history and a negative correlation with sleep rebound, whereas CG9377 shows the opposite pattern, with a positive correlation with rebound sleep and a negative correlation with sleep history. Gene expression values (log₂ TPM) were averaged across three biological replicates, and sleep measurements were averaged across the number of flies per sample.

*CG9377*, a predicted serine-type endopeptidase family member, exhibits the opposite response, i.e., a negative correlation with sleep history and positive correlation with sleep rebound (Fig. 14). *CG9377* is selectively expressed in ensheathing glia and is down regulated by water deprivation^78^. Notably ensheathing glia have been implicated in sleep homeostasis^79^. Taken together, this correlation analysis identifies markers that track sleep-wake history and predict sleep rebound with *bona fide in vivo* sleep functions.

## Discussion

In this study, we sought to identify core molecular mediators of sleep homeostasis that operate independent of the method used to manipulate the sleep–wake cycle. By integrating whole-brain transcriptomics across baseline conditions, mechanical sleep deprivation, thermogenetic and optogenetic manipulations, and pharmacologic sleep induction, we did not identify a universal high-amplitude transcriptional marker of sleep pressure. Instead, sleep homeostasis appears to emerge from distributed and partially overlapping gene programs, some method-specific and others shared across perturbations. Across analytical approaches, including differential gene expression, dataset overlap, correlation with sleep history and rebound, and GPT-assisted literature mining (fl.ai), we identify candidate genes and pathways with functional in vivo relevance, including neuropeptidergic signaling (e.g., *SIFa*, *AstA-R2*), the NA/UNC79/UNC80 sodium leak channel complex, ribosomal components, immune effectors, and mitochondrial oxidative phosphorylation genes. Together, these findings argue that sleep homeostasis is not encoded by a single transcriptional signature but instead reflects convergent, multi-pathway molecular processes that integrate sleep–wake history across time scales and contribute to rebound sleep.

Our data indicate that the transcriptomic response to one method of sleep manipulation shows surprisingly limited overlap with that induced by alternative approaches. Importantly, we did not identify a universal transcriptomic marker of sleep homeostasis across all datasets, even when lowering our threshold to log2FC ≥ 0.5, well below the magnitude observed for core circadian clock genes. One possibility is that such factors may be evident in a cell-type specific manner necessitating more targeted^80^ or higher resolution single cell methods^33, 81^. Nonetheless, these results suggest that sleep homeostasis is not governed by a single large-amplitude transcriptional program, but rather by context-dependent molecular responses.

Despite this heterogeneity, we observed significant overlap between genes modulated under baseline sleep–wake conditions and those altered by multiple sleep perturbations, most notably mechanical sleep deprivation. This pattern supports the idea that homeostatic processes are continuously engaged during normal wakefulness, rather than being triggered solely by extreme perturbation. These findings are consistent with two-process models of sleep regulation as well as forced desynchrony studies ^2, 82, 83^. Indeed, transcriptomic studies in mouse cortex have also found a significant overlap between circadian and sleep deprivation regulated genes^84^. We propose that core homeostatic mechanisms should therefore be evident under baseline conditions.

By integrating transcriptomic data across diverse sleep–wake conditions with GPT-assisted literature mining, differential gene expression analysis, and gene correlation approaches, we identified genes that are consistently modulated across sleep–wake states under multiple conditions and that alter sleep in vivo when genetically perturbed in a manner consistent with a homeostatic function. Among these are *unc79*, a component of the sodium leak channel complex; the neuropeptide SIFa; the glucose transporter *rumpel*; *AstA-R2*, an AstA receptor, and *Trh*, the rate limiting enzyme for serotonin synthesis. Together, these findings highlight the diverse molecular architecture underlying sleep homeostasis, spanning ion channel regulation, neuropeptidergic and neuromodulator signaling, metabolism, and nutrient transport. We also identify a ribosomal protein gene regulated across several sleep–wake manipulations, suggesting a wake-dependent role in ribosome biogenesis and potentially global protein translation. Of note, genetically increasing ribosomal RNA expression promotes sleep^85^. This raises the possibility that sleep–wake history dynamically shapes translational capacity to mediate sleep homeostasis^38, 86^.

An examination of orthologous pathways in mammals suggests that the functional homeostatic genes identified in Drosophila are evolutionarily conserved. The mammalian orthologs of *Drosophila na*, *unc80*, and *unc79*—*NALCN*, *UNC80*, and *UNC79*, respectively—have also been genetically linked to sleep regulation. A large-scale chemical mutagenesis screen in mice identified a semidominant gain-of-function mutation in *Nalcn*, termed *dreamless*, that produced a marked reduction in REM sleep^87^. Notably, this mutation also altered theta-range power during NREM sleep, suggesting a broader role in sleep regulation beyond REM sleep alone. Consistent with these findings, mutations in human *UNC80*^88^ and *NALCN*^89–91^ have also been associated with altered sleep levels or sleep organization.

Our identification of a homeostatic role for the serotonin biosynthetic enzyme tryptophan hydroxylase is likewise consistent with mammalian studies. In both zebrafish and mice, activation of serotonergic raphe neurons promotes sleep, whereas ablation of these neurons impairs both sleep and the homeostatic response to sleep deprivation, supporting a conserved role for serotonergic signaling in sleep homeostasis^92^.

Conservation is also evident at the level of gene regulation. Mitochondrial electron transport genes are upregulated in hippocampal astrocytes following sleep deprivation, although the opposite response has been reported in the prefrontal cortex^93^, suggesting region-specific regulation. In addition, genes involved in protein translation and 40S ribosome assembly are enriched among synaptic transcripts upregulated near the end of the natural wake period (pre-dusk) in mice^31^. This daily modulation is disrupted by sleep deprivation, suggesting that these pathways are sensitive to sleep pressure and may participate in homeostatic regulation. Several transcripts encoding ribosomal proteins are upregulated during wakefulness in rodents, including genes encoding *Rpl21*, *Rpl17*, and *Mrpl23*^94, 95^. E-box upstream elements and bHLH transcription factors have also been implicated in mammalian sleep^96^. Although a more systematic cross-species analysis will be required, these findings collectively suggest that key sleep–wake-dependent pathways and transcriptional programs may be conserved between flies and mammals.

Collectively, our findings do not support the existence of a single universal transcriptional marker of sleep pressure. Rather, sleep homeostasis appears to arise from distributed molecular programs spanning energy metabolism, redox regulation, neuronal excitability, neuropeptidergic signaling, and protein synthesis, consistent with the large number of diverse sleep genes thus far identified in flies^97^. In this framework, sleep need is not encoded by one dominant transcriptional signature but instead emerges from the coordinated activity of multiple interacting pathways that cumulatively register sleep–wake history over time and drive restorative rebound sleep. Interestingly, enrichment of E-boxes in potential transcriptional regulators sequences of wake-induced genes as well as wake-induced bHLH transcription factors that may target those E-boxes suggest potential key regulators that may orchestrate these diverse pathways. We hypothesize that this distributed logic enables sleep homeostasis to integrate many types of waking experience, to integrate that experience over multiple time scales, and to maintain robustness to noise.

## Acknowledgements

This research was supported by NIH R35NS132223 and the Eagles Autism Foundation to R.A., and NIH F32 NS110183 to C.R.

## Methods

### Fly husbandry and stocks

Flies were maintained on standard media composed of sucrose, yeast, agar and molasses in a 12:12 LD cycle between 21-25°C. Most of the fly lines used for sequencing experiments were acquired from the Bloomington Drosophila Stock Center (BDSC; Bloomington, IN) and include *w*^1118^ *iso31* (#5905), UAS-TrpA1 attP16 (#26263), R21H11-Gal4 (#48960), R60F09-Gal4 (#39255), R65H02-Gal4 (#48234), and R85C10-Gal4 (#40424). HC-Gal4 was obtained from Marco Gallio ^64^. ppk-Gal80 was obtained from William Joiner^63^.

### TrpA1-mediated sleep deprivation and sleep promotion

Male flies expressing Gal4 under the control of various promoters were crossed to virgin females containing a UAS-TrpA1 transgene. Progeny were collected and mated at 21°C in a 12hr:12hr LD cycle. Three to ten-day old female flies were loaded into 5 × 65 mm glass behavior tubes containing 5% sucrose-2% agar medium. Fly behavior was recorded using the Drosophila Activity Monitoring System (DAM System) from Trikinetics and analyzed using Sleepmat software^98^. Activity was measured in 1 min bins and sleep was identified as 5 min of inactivity^99, 100^. For all experiments, flies were allowed to acclimate to the tube during the loading day. Following acclimation, the flies were monitored for at least 24 hrs of baseline sleep at 21°C. Afterwards, flies were subjected to various lengths of exposure to 29°C depending on the experiment. For wake-promoting lines, flies were shifted to 29°C from ZT21 to ZT24 (i.e., for 3 hrs at the end of the night until lights on). For the sleep-promoting line R85C10-Gal4, flies were shifted to 29°C from ZT0 to ZT12 (ie. for all 12 hrs of the lights on period). For all RNA-seq experiments, flies were collected immediately at the end of the 29°C temperature shift. Flies were 5-14 days old at the time of collection for RNA-seq. For behavior experiments, flies were returned to 21°C for measurements of sleep activity in the aftermath of TrpA1 activation. Change in sleep during activation was calculated as the difference between sleep during activation and sleep during the equivalent 12 hr period occurring 24 hrs beforehand at baseline. Change in sleep following activation was calculated as the difference between sleep during rebound immediately following activation and sleep during the equivalent 12 hr occurring 24 hrs beforehand at baseline.

### Mechanical sleep deprivation

Age-matched *Iso31* flies were collected and mated at 25°C in a 12hr:12hr LD cycle. Four to six day-old mated female flies were placed in 5 × 65 mm glass behavior tubes containing 5% sucrose-2% agar medium. Tubes were loaded into small DAMS boards which were subsequently secured on a multi-tube vortexer (VWR-2500) fitted with a mounting plate (Trikinetics, Waltham, Massachusetts, USA) inside an incubator maintained at 25 °C in a 12:12 LD cycle. Following acclimatization on the loading day, a baseline period was recorded for at least 24 hrs. Mechanical sleep deprivation (SD) was performed by delivering a 2 second vibration stimulus randomized within consecutive 20 second intervals throughout the night. Behavioral data was recorded throughout the experiment using the DAM System and data was analyzed using SleepMat software^98^. For SD experiments only flies deprived of >90% of baseline sleep at each SD interval were analyzed. Sleep gain was calculated as the difference between sleep during rebound and sleep during the equivalent 12 hr at baseline. For RNA-seq experiments, flies were collected from the incubator at ZT0 immediately after the end of SD. Each SD was timed to end at lights on — thus, 3 hr SD began at ZT21, 6 hr SD began at ZT18, and 12 hr SD began at ZT12. Flies were 7-9 days old at the time of collection for RNA-seq.

### Circadian profiling

Age-matched *Iso31* flies were collected and mated at 25°C in a 12hr:12hr LD cycle. Four to five day-old mated female flies were placed in 5 × 65 mm glass behavior tubes containing 5% sucrose-2% agar medium. Tubes were loaded into small DAMS boards (Trikinetics, Waltham, Massachusetts, USA) inside an incubator maintained at 25 °C in a 12:12 LD cycle. Following acclimatization on the loading day, a baseline period was recorded for at least 24 hrs. Behavioral data was recorded using the DAM System and data was analyzed using SleepMat software^98^. For RNA-seq experiments, flies were collected from the incubator at ZT0, 4, 8, 12, 16, and 20 for immediate dissection. Flies collected during the dark period (ZT12, 16, 20) were kept out of the light until it was time to dissect. Flies were 6-8 days old at the time of collection for RNA-seq.

### Dissection for whole brain RNA-seq

For whole brain bulk RNA-sequencing experiments, the appropriate flies were collected from incubators as described above. Fly behavior was quickly analyzed using Sleepmat software to rule out dead flies or flies that did not meet the criteria for sleep deprivation (>90% deprivation) or promotion (>90% sleep) depending on the experiment ^98^. For TrpA1 experiments, we collected flies from the experimental group, which contained both a Gal4 and UAS-TrpA1 and were exposed to the temperature pulse, as well as two groups of control flies. One control contained both a Gal4 and UAS-TrpA1 but was maintained at 21°C throughout the experiment while the other contained only a Gal4 and was exposed to the 29°C temperature pulse. For mechanical sleep deprivation and circadian profiling experiments, flies were maintained at a constant temperature (25°C) and kept in a 12h:12h LD cycle. Any flies collected during the dark period of the cycle were kept in the dark until they could be anesthetized with carbon dioxide and decapitated.

Brains were dissected in in ice-cold dissecting saline containing 9.9 mM HEPES-KOH buffer, 137 mM NaCl, 5.4 mM KCl, 0.17 mM NaH2PO4, 0.22 mM KH2PO4, 3.3 mM glucose, 43.8 mM sucrose, pH 7.4 and a cocktail of channel blockers (0.1 μM tetrodotoxin (TTX), 20 μM 6,7-dinitroquinoxaline-2,3- dione (DNQX), and 50 μM D(–)–2-amino-5-phosphonopentanoic acid (AP-5) to block neuronal activity during dissection and processing. Channel blockers were added to the dissecting saline immediately before use. Fifteen brains were used for each replicate and three replicates were used for each condition. Once all brains were dissected, they were transferred to a LoBind microcentrifuge tube and spun down at 2000 rpm for 1 min before removing the dissecting saline. They were washed twice with 500 μL of dissecting saline with TTX, AP5, and DNQX. All the saline was removed and 100 μL of extraction buffer from the Arcturus PicoPure kit was added. Then the brains were vortexed for 30 seconds and incubated at 42°C for 30 min before being stored at -80°C.

### Library Preparation

Once all samples were collected, total RNA was isolated using the Arcturus PicoPure kit with on-column DNaseI digestion. RNA was subjected to quality control assessment on a Bioanalyzer Pico chip and quantified on a Qubit fluorometer. RNA was used to generate sequencing libraries using the NEBNext Ultra II RNA Library Prep Kit and following manufacturer instructions. mRNA was selected prior to reverse transcription using the NEBNext Poly(A) mRNA Magnetic Isolation Module. Library quality was assessed on a Bioanalyzer High Sensitivity DNA chip and quantity was assessed on a Qubit fluorometer. Libraries were sent to Novogene for sequencing on a NovaSeq. At least ten million 150 bp paired end reads were generated per sample.

### RNA Sequencing Data Analysis

Reads were pseudo aligned and quantified using Kallisto (v0.46.1)^101^ against a prebuilt index file constructed from Ensembl reference transcriptomes (v96). Kallisto was used to process paired end reads with 10 bootstraps. Differential expression analysis of the resulting abundance estimate data was then performed with Sleuth (v0.30.0^102^). Gene-level abundance estimates were computed by summing transcripts per million (TPM) estimates for transcripts for each gene. To measure the effect of a particular condition against another condition for a variable, sleuth uses a Wald test which generates p values as well as q values (an adjusted p value using the Benjamini-Hochberg procedure). For rhythmic analysis of circadian profiling control data, circadian samples were collected at six time points: ZT0, ZT4, ZT8, ZT12, ZT16, and ZT20. Rhythmicity was assessed using the RAIN algorithm^103^ with three biological replicates per time point. Genes were considered rhythmic if they met the criteria of log₂(fold change) > 0.5 and Benjamini–Hochberg adjusted Q value (BH Q) < 0.05. Fold change was calculated as the difference between peak and trough expression values (maximum minus minimum expression). RNA-sequencing data generated and analyzed in this study have been deposited in the Gene Expression Omnibus (GEO) under accession number GSE322542.

### Classification of consistently sleep and wake associated transcripts

We analyzed differentially expressed and cycling transcripts from seven whole-brain sleep/wake experiments (Baseline, R85C10>TrpA1, MechSD3, MechSD6, MechSD12, and *THIP, and *R23E10>ChR) to identify genes consistently associated with sleep or wake state (*^41^). Within each experiment, genes were classified by expression direction relative to the manipulation: genes upregulated under sleep-promoting conditions or downregulated under wake-promoting conditions were labeled sleep-associated, whereas the converse were labeled wake-associated. We first defined a higher-amplitude gene set (FC1) using a threshold of |log2FC| > 1 and q < 0.05 as a statistical threshold for classifying a gene as ‘sleep’, ‘wake’ or ‘cycling’ within an experiment. A gene was retained in the consistent sleep set if it was classified as sleep-associated in at least two of the seven experiments and never classified as wake-associated in any experiment; consistent wake genes were defined analogously. Cycling transcripts in baseline, after gating by fold-change and q-value thresholds, were treated as a wild-card supporting both wake- and sleep classifications.

### Overlap analysis

To assess similarity between experiments, we performed pairwise overlap analyses using one-sided Fisher’s exact tests, with the background defined as the intersection of genes detected in both datasets (>50% of samples non-zero within each experiment). P-values were adjusted separately for sleep and wake comparisons using the Benjamini–Hochberg method. We then repeated the same analysis using a lower fold-change threshold (FC0.5; |log2FC| > 0.5, q < 0.05). The sleep-wake analyses, including overlap-enrichment tables and UpSet plots, were generated using a custom Hugging Face Streamlit app. https://huggingface.co/spaces/aadish98/Allada_GeneRef_Search

### Gene Ontology Enrichment Analysis

Functional enrichment analysis was performed using the NIH DAVID web service (DAVID Bioinformatics Resources). Gene lists were submitted as FlyBase gene IDs (FLYBASE_GENE_ID) for Drosophila melanogaster (species 7227). Enrichment was evaluated in GOTERM_BP_DIRECT, GOTERM_CC_DIRECT, GOTERM_MF_DIRECT, and KEGG_PATHWAY categories using DAVID Functional Annotation Chart (minimum term count = 1). Terms with FDR < 10% were defined as statistically significant. The CLI tool used for executing the analysis and visualization plots are published on GitHub: https://github.com/aadish98/GO_Analysis

### Motif Enrichment Analysis

Motif enrichment analysis was performed using HOMER (Hypergeometric Optimization of Motif EnRichment)^52^. Gene lists corresponding to sleep and wake associated genes (log_2_FC > 0.5) were analyzed using the findMotifs.pl function with the *Drosophila melanogaster* genome as reference. Promoter regions were defined as −1000 bp upstream to +50 bp downstream of the transcription start site (TSS). HOMER automatically generated background sequences matched for GC content and sequence composition. Enriched transcription factor binding motifs were identified using a hypergeometric test, and motifs were ranked based on statistical significance (p-value). The top enriched motifs were selected for visualization.

### Correlation Between Sleep History or Rebound and Gene Expression

Correlation analyses were performed using gene-level TPM values. Genes with TPM < 1 in more than 50% of samples were excluded to remove low-expressed genes that could produce unstable or spurious correlations. For each experimental condition, the three biological replicates were averaged, and the resulting mean TPM values were log-transformed prior to correlation analysis. Behavioral sleep measurements were averaged across all flies within each condition. We examined the relationship between gene expressions and two types of sleep metrics:

1. Sleep before sampling (sleep history) Cumulative sleep was calculated over increasing time windows prior to sample collection (1 h, 2 h, …, up to 12 h before sampling). For example, the 2-hour value represents the total sleep across the preceding 2 hours, not a rolling 1-hour window. Pearson correlations were computed between log-transformed gene expressions and each of these 12 cumulative sleep intervals. All samples were included in this analysis.
2. Sleep rebound Sleep rebound was defined as the difference between cumulative sleep after sampling (1–12 h) and the corresponding cumulative baseline sleep (1–12 h) for each condition. Pearson correlations were computed between gene expression and each of the 12 rebound intervals. Circadian-control samples were excluded from this analysis.

All correlations were assessed using Pearson correlation coefficients. Statistical significance was determined using Benjamini–Hochberg–adjusted p-values, with q < 0.05 considered significantly correlated.

### LLM-Assisted Literature Review and Gene Classification for Sleep/Circadian Genes

To systematically evaluate whether candidate sleep-wake associated and correlated genes have published evidence linking them to sleep or circadian biology, we developed an LLM-assisted literature review pipeline (fl.AI-CLI). The pipeline converts input gene symbols to FlyBase identifiers, then retrieves candidate references from three complementary sources (FlyBase, PubMed, Europe PMC), deduplicates them at the PMCID level, and ranks by cross-source agreement and recency. References are filtered by keyword relevance (sleep, circadian) in title/abstract metadata, and full text is retrieved where available via open-access repositories (PMC Open Access, Europe PMC, PMC HTML, Unpaywall, OpenAlex, Crossref and DOI resolver.) For each gene, an LLM (GPT5-nano) summarizes gene-specific evidence (functional role of gene, supporting phenotypic evidence and reagents used to illicit the phenotype) from up to six passing and relevant references; reference summaries are aggregated per gene, which are then classified by an LLM (GPT5-mini) into keyword categories “sleep”, “circadian” (or both) with a confidence score (0–100) and brief rationale. The output is an annotated spreadsheet containing per-gene classifications and the supporting reference summaries used to derive them. The CLI tool is made publicly available through GitHub: https://github.com/aadish98/fl.AI-gene-classification.

### Declaration of generative AI and AI-assisted technologies in the manuscript preparation process

During the preparation of this work, the authors used ChatGPT for editing and grammar of specific sections. The authors reviewed and edited the output as needed and take full responsibility for the content of the published article.

---

**Figure S1.**
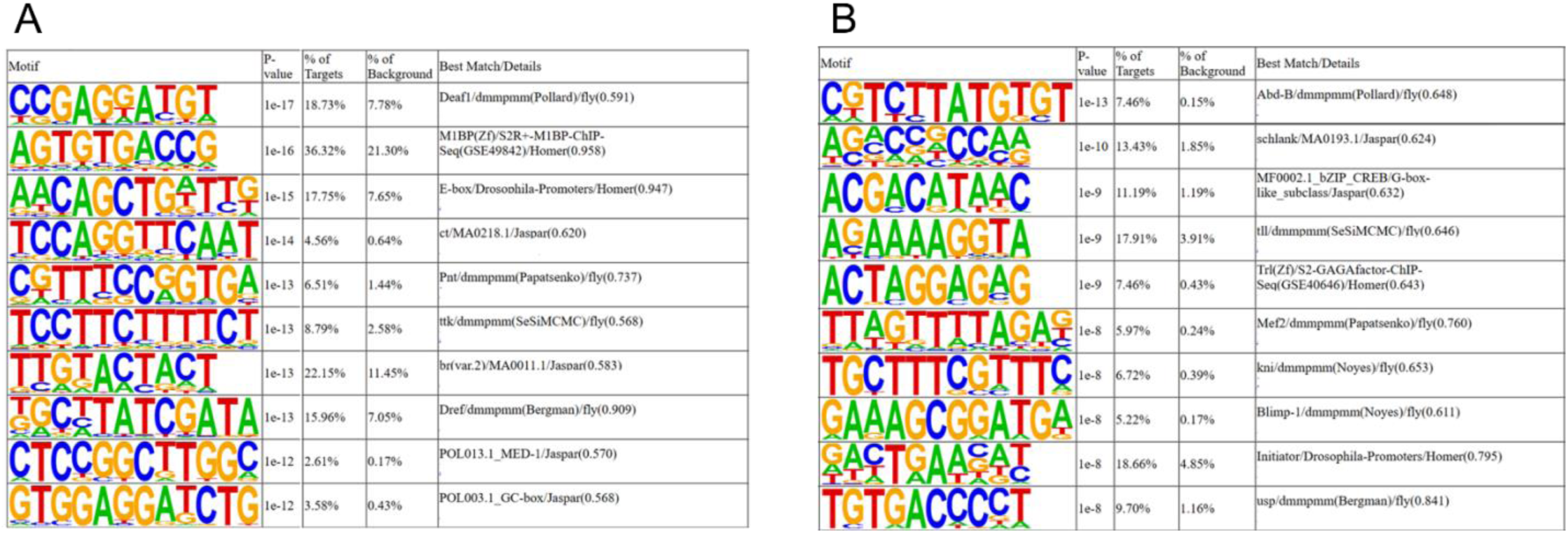
Motif enrichment analysis of wake and sleep conditions. (A) Top 10 transcription factor binding motifs enriched in promoter regions of wake-associated genes identified using HOMER. (B) Top 10 motifs enriched in promoter regions of sleep-associated genes. Motifs are ranked based on enrichment p-value. For each motif, the p-value, percentage of target sequences, percentage of background sequences, and best match are shown.

**Figure S2.**
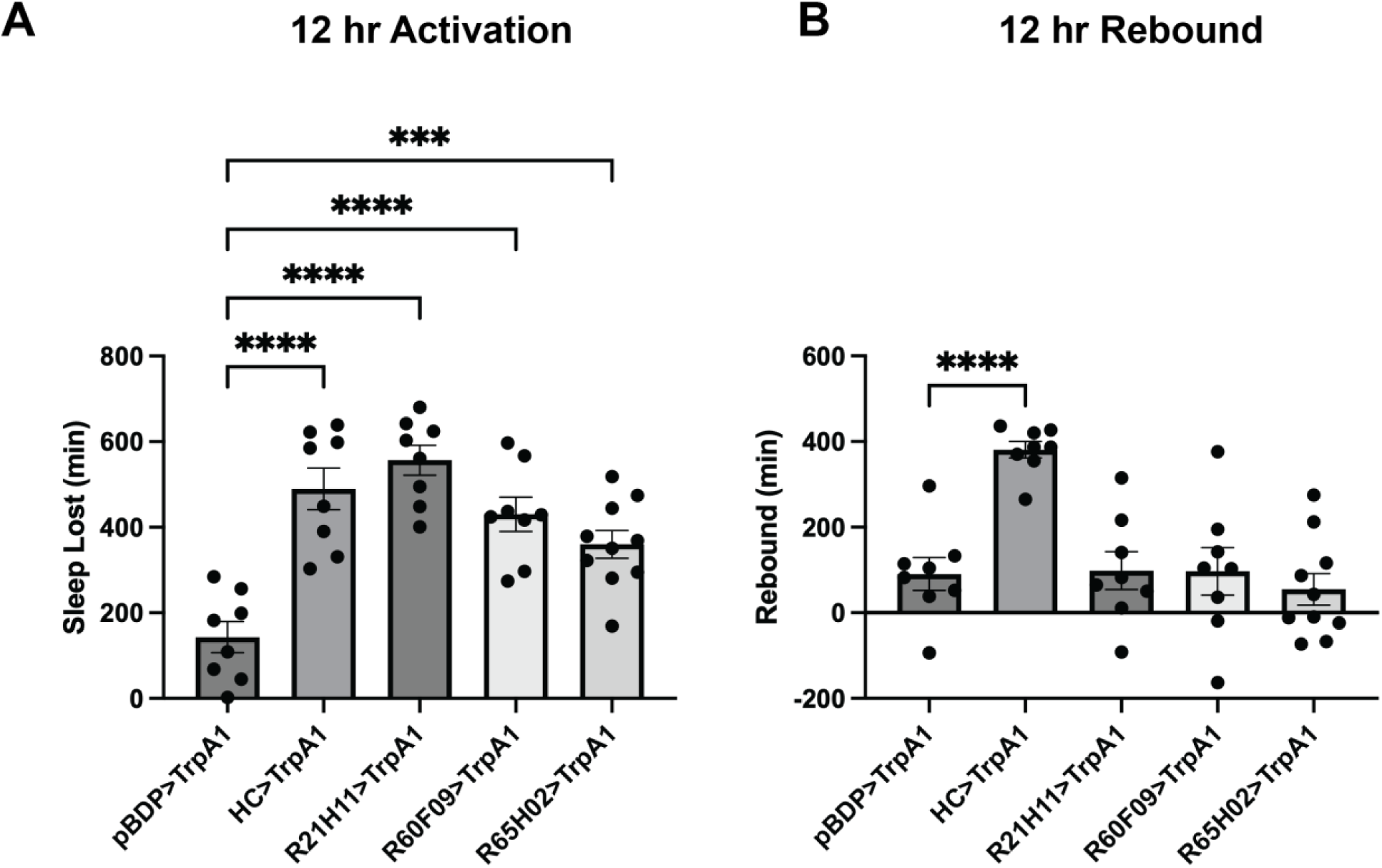
Wake Promotion and Rebound Sleep after 12h TrpA1 Activation. GAL4 lines for sleep deprivation were chosen following 12 hr activation experiments. A. Sleep lost during 12 hr nighttime activation of GAL4 lines at 29°C compared to the 12 hr period occurring one day before activation. Outside of activation, flies were maintained at 21°C in a 12:12 LD cycle. Each GAL4 was compared to an enhancer-less GAL4 driver (pBDP-GAL4). *** p = 0.0008, **** p<0.0001 by ANOVA with Dunnett’s multiple comparison test. B. Rebound sleep during the 12 hr period immediately after activation of GAL4 lines. Each GAL4 was compared to an enhancer-less GAL4 driver (pBDP-GAL4). **** p<0.0001 by ANOVA with Dunnett’s multiple comparison test.

**Figure S3.**
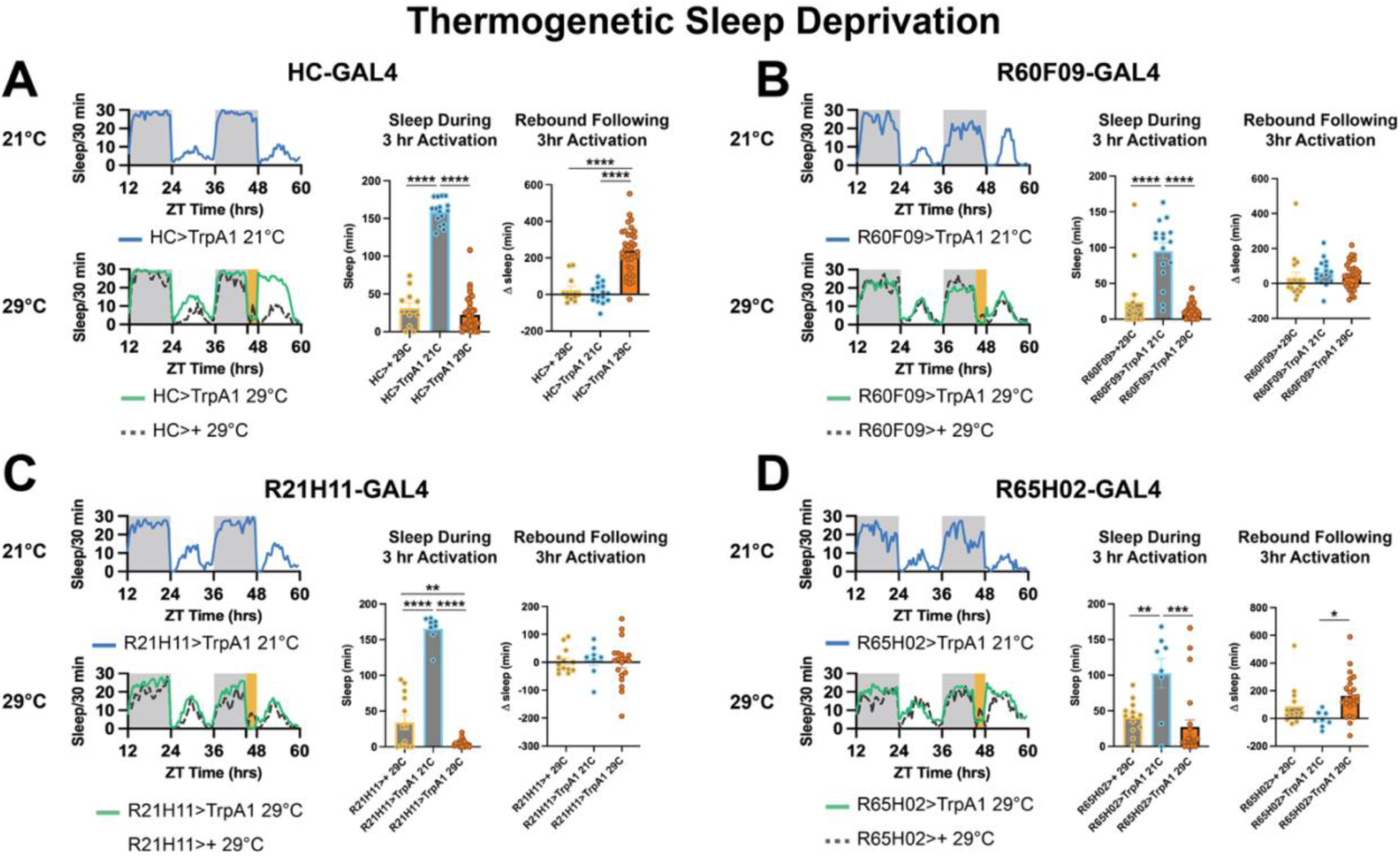
Wake promotion via 3 hour thermogenetic TrpA1 activation. A-D. Panels A through D show four GAL4 lines that were used to induce sleep deprivation when paired with TrpA1 and a temperature pulse to activate neurons expressing TrpA1. The top left graph in each panel shows two circadian cycles of sleep behavior at 21°C depicting flies bearing both the GAL4 and UAS-TrpA1 transgenes. The bottom left graph in each panel shows two circadian cycles of sleep behavior for two genotypes: a GAL4 line paired with a UAS-TrpA1 transgene as well as a GAL4/+ line. The flies are maintained at 21°C for most of the experiment except for a brief 3 hr transition to 29°C at the end of the second night (depicted in orange). The two other graphs in each panel show a quantification of total sleep during the 3 hr activation for each condition (left) and the change in sleep during the 12 hr rebound period following activation compared to the previous day (right). (A) Activation of HC-GAL4 neurons promotes wakefulness and over 200 minutes of rebound sleep following activation. HC>TrpA1 21°C (n = 16), HC>+ 29°C (n = 11), HC>TrpA1 29°C (n = 35). **** p<0.0001 by ANOVA with Tukey’s multiple comparisons. (B) Activation of R60F09-GAL4 neurons promotes wakefulness without rebound sleep following activation. R60F09>TrpA1 21°C (n = 16), R60F09>+ 29°C (n = 16), R60F09>TrpA1 29°C (n = 32). **** p<0.0001 by ANOVA with Tukey’s multiple comparisons. (C) Activation of R21H11-GAL4 neurons promotes wakefulness without rebound sleep following activation. R21H11>TrpA1 21°C (n = 8), R21H11>+ 29°C (n = 13), R21H11>TrpA1 29°C (n = 19). ** p = 0.0039, **** p<0.0001 by ANOVA with Tukey’s multiple comparisons. (D) Activation of R65H02-GAL4 neurons promotes wakefulness with limited rebound sleep following activation. R65H02>TrpA1 21°C (n = 8), R65H02>+ 29°C (n = 13), R65H02>TrpA1 29°C (n = 23). * p = 0.0146, ** p = 0.0097, **** p = 0.0005 by ANOVA with Tukey’s multiple comparisons.

**Figure S4.**
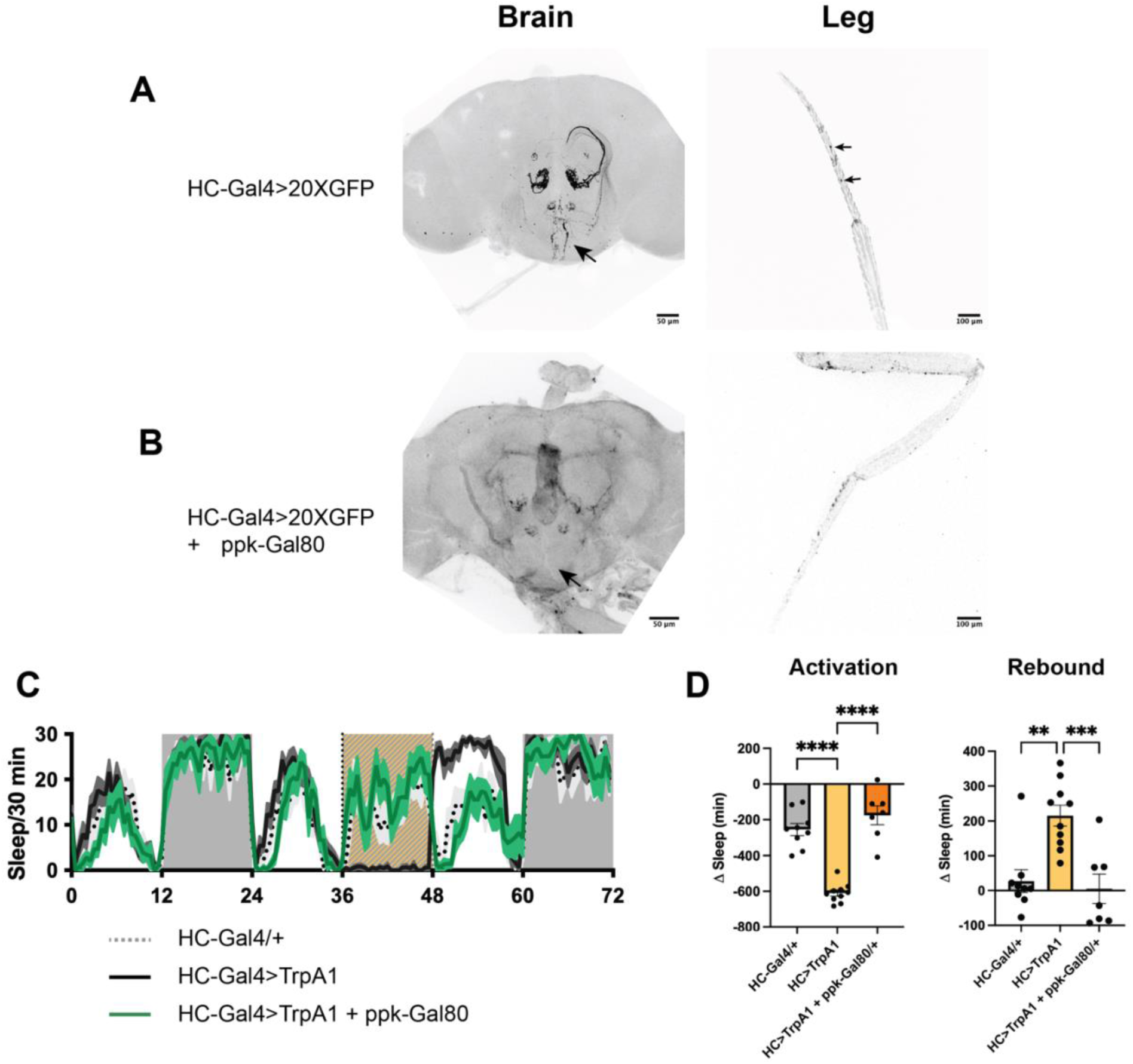
*pickpocket* leg neurons are necessary for HCGAL4 wake promotion and rebound. Wake promotion and rebound in HC-GAL4 experiments is driven by ppk+ cells in the legs. A. Confocal imaging of the brain and leg of flies in which HC-GAL4 was used to drive expression of GFP. GFP is shown in black. Arrow on the brain indicates terminals of the leg neurons in the subesophageal zone (SEZ) of the brain previously reported in Seidner et al. Arrows on the leg indicate two cell bodies associated with wake-promotion {Satterfield, 2022}. B. Confocal imaging of the brain and leg of flies in which HC-GAL4 was used to drive expression of GFP in the presence of a ppk-Gal80 transgene blocking expression in ppk neurons. The absence of terminals in the SEZ is indicated by the arrow. Cell bodies in the distal part of the leg are also missing. C. Coexpression of ppk-Gal80 completely blocked wake promotion during a 29°C nighttime pulse indicated in orange. D. Quantification of the data in panel C indicates that coexpression of ppk-Gal80 abrogated the wake promoting effects observed when activating HC-GAL4 neurons with a TrpA1 transgene. **** p<0.0001 by ANOVA with Tukey’s multiple comparisons. E. Quantification of the data in panel C indicates that coexpression of ppk-Gal80 abrogated the wake promoting effects observed when activating HC-GAL4 neurons with a TrpA1 transgene. ** p = 0.0013, *** p = 0.0008 by ANOVA with Tukey’s multiple comparisons.

